# Characterization of bacterial intrinsic transcription terminators identified with TERMITe – a novel method for comprehensive analysis of Term-seq data

**DOI:** 10.1101/2024.05.16.594410

**Authors:** Jan Grzegorz Kosiński, Sandeepani Ranaweera, Agnieszka Chełkowska-Pauszek, Mikhail Kashlev, Paul Babitzke, Marek Żywicki

## Abstract

In recent years, Term-seq became a standard experimental approach for high-throughput identification of 3’ ends of bacterial transcripts. It was widely adopted to study transcription termination events and 3’ maturation of bacterial RNAs. Despite widespread utilization, a universal bioinformatics toolkit for comprehensive analysis of Term-seq sequencing data is still lacking. Here, we describe *TERMITe*, a novel method for the identification of stable 3’ RNA ends based on bacterial Term-seq data. *TERMITe* works with data obtained from both currently available Term-seq protocols and provides robust identification of the 3’ RNA termini. Unique features of *TERMITe* include the calculation of the transcription termination efficiency using matched RNA-seq data and the comprehensive annotation of the identified 3’ RNA ends, allowing functional analysis of the results. We have applied *TERMITe* to the comparative analysis of experimentally validated intrinsic terminators spanning different species across the bacterial domain of life, revealing substantial differences in their sequence and secondary structure. We also provide a complete atlas of experimentally validated intrinsic transcription termination sites for 13 bacterial species, including *Escherichia coli, Bacillus subtilis, Listeria monocytogenes, Enterococcus faecalis, Synechocystis sp*., *Streptomyces clavuligerus, Streptomyces griseus, Streptomyces coelicolor, Streptomyces avermitilis, Streptomyces lividans, Streptomyces tsukubaensis, Streptomyces venezuelae*, and *Zymomonas mobilis*.

## INTRODUCTION

Transcription termination in bacteria has been historically documented to occur via two mutually exclusive mechanisms triggered by features in the nascent RNA (intrinsic termination) or by the termination factor Rho (Rho-dependent termination) (1). A canonical intrinsic terminator, or Rho-independent terminator, consists of a G/C-rich RNA hairpin immediately followed by a uridine-rich tract of 8-10 nt (2, 3). Working together, the weak rU-dA hybrid and the adjacent termination hairpin undermine the extraordinary stability of the elongation complex at intrinsic terminator sites (4). Intrinsic termination has traditionally been described as occurring without the participation of any additional regulatory factors (5). However, in recent years intrinsic terminators have been shown to require NusA, NusG, and/or Rho to facilitate efficient termination, indicating that they are not factor independent (6–8).

Rho is an ATP-hydrolyzing RNA helicase that stimulates termination of paused RNA polymerase (RNAP) after binding to an unstructured C-rich and G-poor sequence approximately 70-80 nt in length with regularly interspersed dipyrimidine repeats called a *rut* (Rho utilization) site (9). Traditional models of Rho-dependent termination postulated that after loading on a *rut* site, Rho translocates along the transcript using its ATP hydrolysis activity until it reaches paused RNAP and stimulates transcript release (1). However, according to recent structural studies, before interacting with RNA, Rho can form a pretermination complex (PTC) with RNAP and the elongation factors NusA and NusG, which stabilize the PTC (10, 11). After recruitment, Rho contacts a *rut* site due to several rearrangements of NusA, RNAP, upstream DNA, and Rho. The interaction with *rut* is necessary for the stimulation of Rho ATPase activity, which induces displacement of NusG and weakening of RNAP’s grip on DNA and RNA, resulting in RNAP inactivation and transcript release (11). A recent structural study implies that ATP-dependent translocation by Rho exerts a mechanical force on RNAP leading to transcript release (12).

Post-transcriptional events, such as RNA decay, contribute to the regulation of gene expression in response to changing environmental conditions. mRNA decay is generally initiated by an endoribonucleolytic cleavage by RNase E in *E. coli* and RNase Y in *B. subtilis* (13). The cleaved transcripts are then subject to exoribonucleolytic decay in the 3’-5’ and/or 5’-3’ directions. RNA secondary structures can affect RNase activity in different ways (13). In particular, the strong RNA hairpins associated with intrinsic terminators can serve as a barrier to 3’-5’ exoribonucleases, leading to the generation of transcripts with truncated 3’ ends, thus masking the authentic 3’ ends generated by intrinsic termination.

Term-seq is a high-throughput method allowing genome-wide identification of 3’ RNA ends which is particularly useful in research focusing on transcription termination and RNA processing in bacteria. Two distinct variations of Term-seq were published in (8, 14) (Figure 1). The original protocol by Mondal et al. involves the ligation of a blocked 2’,3’-dideoxy RNA oligonucleotide to rRNA-depleted RNA, followed by a standard bacterial RNA-seq protocol, including random shearing and reverse transcription using random hexamers (8). The resulting cDNA library, which is sequenced using the Illumina platform, along with Term-seq reads, also yields standard RNA-seq reads. Thus, data analysis requires separating the reads into two datasets: all reads (RNA-seq dataset) and reads found to contain the ligated RNA oligonucleotide (Term-seq dataset). After removing the sequence corresponding to the RNA oligonucleotide, Term-seq reads can be used to establish 3’ RNA termini as 5’ read ends reflect 3’ transcript termini. The RNA-seq reads can be used for gene expression analysis or assessment of termination efficiency at identified termination sites (8).

**Figure 1.**
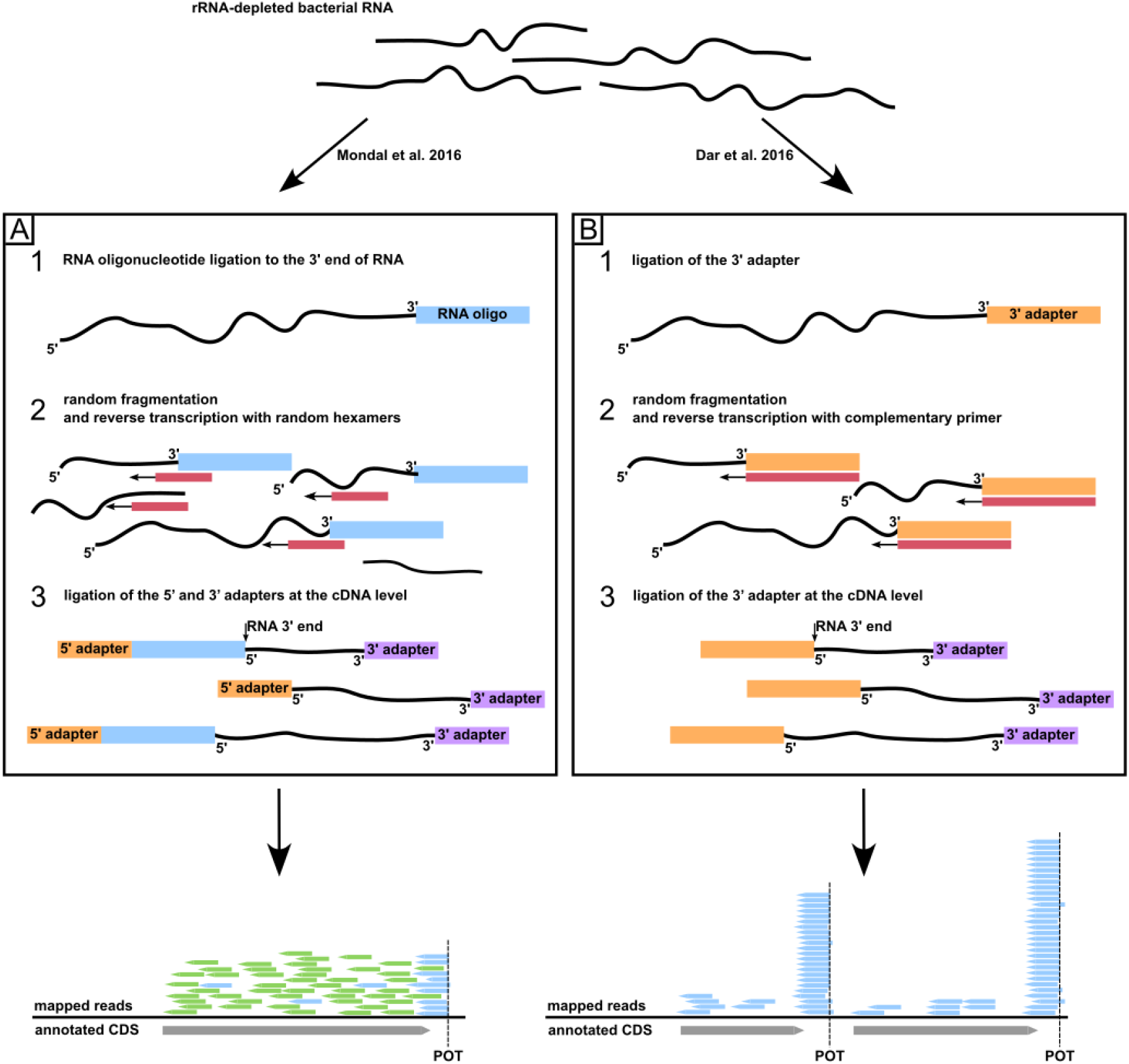
Schematic representation of the Term-seq protocol. a) Term-seq method by Mondal et al. (8). First, a 2’,3’-dideoxy RNA oligonucleotide of known sequence is ligated to the 3’ RNA ends of rRNA-depleted samples (pt. 1). The RNA pool is then subjected to random shearing (pt. 2), reverse transcription using random hexamers as priming sequences (pt. 2), and ligation of the 5’ and 3’ adapters at the cDNA level (pt. 3). After sequencing and trimming of the oligonucleotide sequence, all reads represent a standard RNA-seq library, while the subset that contained the oligonucleotide sequence (Term-seq reads, blue) can be used to reveal the 3’ RNA termini. b) The Term-seq protocol published by Dar et al. (14). First, a 3’ adapter is ligated to the 3’ RNA termini (pt. 1) followed by random fragmentation of RNA (pt. 2), and reverse transcription using primers complementary to the 3’ adapter (pt. 2). The 3’ DNA adapter is ligated to the first strand cDNA (pt. 3). After sequencing, the 5’ ends of the reads point to the 3’ RNA termini.

The protocol by Dar et al. (14) involves the ligation of a 3’ RNA adapter to rRNA-depleted bacterial RNA, random fragmentation of RNAs, reverse transcription primed by a specific oligonucleotide complementary to the ligated 3’ adapter, and ligation of the second adapter at the cDNA level. The method is similar to Mondal et al. in the way that the 5’ ends of obtained sequencing reads correspond to the 3’ transcript termini. The main difference is the selective incorporation into the cDNA library of fragments derived from transcript ends due to the initiation of reverse transcription using a primer complementary to the oligonucleotide ligated to the 3’ ends of RNAs. On the one hand, the resulting dataset provides superior coverage of transcript 3’ termini, but on the other, it does not provide the RNA-seq reads, hampering the simultaneous analysis of gene expression and transcription termination efficiency.

Term-seq has been successfully applied to multiple bacterial and archaeal species, including *B. subtilis* (6–8, 14, 15), *Listeria monocytogenes* (14), *Enterococcus faecalis* (14), *Synechocystis sp*. (16), *E. coli* (17), *Streptomyces sp*. (18–22), *Pseudomonas aeruginosa* (23), *Zymomonas mobilis* (24), *Methanosarcina mazei*, and *Sulfolobus acidocaldarius* (25). Most studies rely on in-house custom analytical approaches to identify and analyze either RNA decay products or transcription termination sites. However, substantially increasing utility of the Term-seq protocols requires efficient algorithms and pipelines that can be used for the identification and analysis of stable 3’ RNA ends. The main challenge is to distinguish genomic locations of stable 3’ ends of transcripts from regions that are highly inconsistent between replicates and may correspond to intermediates of RNA decay, posttranscriptional processing products, incorrectly mapped reads, sequencing artifacts, or simply 3’ ends of poorly expressed transcripts. The only tool currently available for such analysis is *termseq-peaks* (26) *(https://github.com/nichd-bspc/termseq-peaks*), which has been designed and tested to work with Term-seq data from replicated experiments prepared with the Dar et al. protocol. The workflow includes identification of peaks in genomic coverage by 5’ ends of term-seq reads in each replicate, calculation of the Irreproducible Discovery Rates (IDR) between replicates for identified peaks, estimation of the minimal number *N* of peaks below the specified IDR threshold among replicate comparisons, second peak calling on coverage merged from all replicates and selection of *N* peaks with highest prominence. However, this approach is prone to biases caused by highly expressed genes and highly abundant degradation products. Furthermore, *term-seq peaks* do not provide any annotation of the genomic context on the identified 3’ RNA ends.

Here we present *TERMITe*, which is software for identification and annotation of stable 3’ RNA ends based on bacterial TERM-seq data, which overcomes the issues mentioned above. Our method implements a modified version of a *Term-seq peaks* algorithm that overcomes its expression bias by selecting the most probable 3’ RNA ends based solely on reproducibility between replicates. It is designed to work with data derived from both published Term-seq protocols, including utilization of RNA-seq reads obtained with the protocol by Mondal et al. for calculation of transcription termination efficiency. Furthermore, *TERMITe* provides annotation of the results with information that supports the assessment of the biological origin and genomic context of the identified 3’ RNA ends, being the first bioinformatic tool that allows a comprehensive analysis of Term-seq data.

## MATERIAL AND METHODS

### *TERMITe* algorithm

*TERMITe* was designed to work with data obtained from at least two biological replicates of TERM-seq experiment, however, three or more are recommended. It enables the identification of statistically reproducible 3’ RNA termini peaks in genomic coverage by 5’ ends of Term-seq reads. The input coverage data should be calculated independently for each replicate, normalized to sample sequencing depth, and provided in the BIGWIG format for each genomic strand separately. This process can be easily achieved using *bamCoverage* from the deepTools package (27). Optionally, the user can also provide data with genomic coverage from the corresponding RNA-seq reads obtained from each sample, allowing calculation of the termination efficiency for each site (6–8).

*TERMITe* consists of two modules that are designed to be executed consecutively. The first module (*termite find_stable_rna_ends*) is responsible for the identification of stable 3’ RNA ends, while the second module (*termite annotate*) is responsible for the annotation of the termination sites. The algorithm for identifying stable 3’ RNA ends is based on the work of Adams et al. (*https://github.com/NICHD-BSPC/termseq-peaks*) with several adjustments and modifications. First, by analysis of genomic coverage by 5’ ends of the Term-seq reads (corresponding to 3’ ends of RNA), the coverage peaks are identified using the *find_peaks* function from the Python module *scipy*.*signal* (28). This process is repeated independently for each replicate and strand with relaxed (default) settings to enable the identification of all possible peaks. Then, positions at both ends of each peak with coverage lower than 10% of total peak coverage are trimmed. The identified peaks are then subjected to cross-replicate comparison to identify reproducible termination events, which is achieved by calculating Irreproducible Discovery Rates (IDR) (version 2.0.3; https://github.com/nboley/idr). IDR was proposed as an excellent method to distinguish reproducible Term-seq peaks from random background noise in *termseq-peaks* (26). In *TERMITe*, an IDR value is calculated for each pair of overlapping peaks across two replicates. Peaks with an IDR value lower than a specified threshold (0.05 by default) in at least M pairwise comparisons of replicates (1 by default) are extracted and reported as potential 3’ RNA termini. The reported position of the 3’ RNA end corresponds to the peak summit, which is the highest point of the peak across all the replicates. In the case of multiple overlapping positions with the same maximum height, the most downstream position is reported. Optionally, *TERMITe* calculates the termination efficiency for each identified 3’ RNA end. This value enables the analysis of intrinsic transcription termination dynamics or nuclease processing efficiency and should be interpreted as the ratio of transcript isoforms ending at any given 3’ RNA termini to longer transcripts. The calculation is based on the genomic coverage of the termination region by RNA-seq reads derived from all provided biological replicates (normalized to sequencing depth). Coverage in a window of 13 nt upstream and 13 nt downstream from the peak summit is extracted, in both cases excluding three positions closest to the termination site. The termination efficiency (*T*) is calculated as

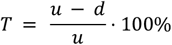

where *u* is the average coverage from the remaining 10 nt upstream from the peak summit and *d* 10 nt downstream. To provide a reliable estimate of termination efficiency and exclude regions with locally disturbed RNA-seq coverage, the calculation is performed only if the average upstream coverage is higher than a particular threshold (RPM 0.25 by default). Stable 3’ RNA ends identified with this module are saved in a tabular *bedNarrowPeak* format.

Since *TERMITe* works by analysis of 5’ ends of Term-seq reads, the identified set of 3’ RNA ends is composed of transcription termination sites and 3’ ends of stable RNA degradation intermediates, which are in most cases indistinguishable from termination sites. The number of nuclease-derived sites depends on the presence of the 3-OH group in the RNA degradation/cleavage products and their susceptibility to ligation of the 3’adapter during the Term-seq procedure.

The second module, *TERMITe annotate*, was developed to provide a biological context of the stable 3’ RNA ends identified. It utilizes the provided genomic annotation to assign genomic features to POTs and predicts whether the observed transcript end is a likely direct outcome of intrinsic transcription termination. *TERMITe annotate* also allows the comparison of stable 3’ RNA ends identified for numerous samples representing multiple experimental conditions. The output of this step can be used directly for the differential termination analysis, e.g., based on termination efficiencies.

First, the algorithm identifies overlapping peaks from different samples using *bedtools merge* (29, 30) with the interface of *pybedtools* (31). Reference Points of Observed [RNA] Termini (POT) are calculated based on the mean genomic coverage by the overlapping 3’ RNA ends across all samples and replicates. The POT corresponds to the position with the highest coverage. The most downstream position is reported in the case of ties. When gene annotations are provided, *TERMITe* reports potential overlapping and closest genes, both upstream and downstream, with respective distances to the POT. Additionally, the report includes feature types from the provided GTF/GFF3 files (representing gene annotations) overlapping the 3’ termini (e.g., CDS, tRNA, 3’ UTR, 5’ UTR, etc.). This step is performed using the Python package *gffutils* (31). Additionally, *TERMITe* can report overlaps between *TERMITe* peaks and any number of custom features described by genomic coordinates in the form of one or more BED files. This report can be particularly useful for annotating the results with prior knowledge (e.g. known terminators from reference databases) or results of other experiments.

*TERMITe* implements two methods for examining the possibility of forming the intrinsic terminator hairpin upstream of the POT. The first one is based on a well-established method *TransTermHP* (32) designed to predict intrinsic transcription terminators in a genome-wide manner. *TERMITe* assigns terminators detected by *TransTermHP* to the identified stable 3’ RNA ends (maximum and minimum distances between the upstream terminator hairpin and the stable 3’ RNA end can be set using appropriate parameters). The calculated distance between the hairpin and POT is reported, along with the typical terminator features, such as potential A tract upstream of the hairpin, downstream U tract, hairpin sequence and structure as predicted by the *TransTermHP*. The position of the POT is marked with the capital letter within the sequence, making it easy to locate in the results. Whenever the distance from the POT to the base of the predicted terminator hairpin is within the specified range ([0 to 10] by default), the 3’ RNA end is reported to be the product of intrinsic termination. This protocol is called the *TransTermHP protocol*.

To increase the sensitivity of detection of less conserved intrinsic terminators in POT regions, we developed a second method, which is based on scanning of the sequence from 80 nt upstream to 30 nt downstream from the POT with *TransTermHP* using a permissive set of parameters to maximally increase the method’s sensitivity *(--all-context, --min-conf=0, --min-stem=4, --min-loop=4, --max-len=40, --max-loop=20, --uwin-require=1, --uwin-size=6*). The most probable terminator is selected based on hairpin confidence, hairpin score, and tail score (criteria are evaluated in this specific order). The sequences of identified hairpins are next folded using *RNAfold* (33), and the free energy of the hairpin structure is evaluated (required to be ≤ -3 kcal/mol). The distance between the hairpin and the POT must be within the range specified by the user to report the stable 3’ RNA as an intrinsic termination site ([0 to 10] by default). This protocol is called the *RNAfold protocol*.

The *TERMITe* annotation output is saved in a tabular file (tab-separated values). A detailed description of the tool usage, required and optional parameters, the structure and contents of the output files, as well as the example test case, are available at https://github.com/zywicki-lab/TERMITe.

### Library preparation and sequencing

Three biological replicates of the wild-type strain of *E. coli* (NB1246) were grown in LB medium supplemented with 25 μg/mL kanamycin at 37 °C. Cells were collected during the mid-exponential growth phase, and total RNA was extracted using the RNeasy mini kit (Qiagen). The RNA was then dephosphorylated by treatment with calf intestinal alkaline phosphatase (CIP) and the rRNA was depleted using the MICROBExpress rRNA depletion kit (Thermo Fisher Scientific). A unique 2’,3’-dideoxy RNA oligonucleotide (IDT) that was phosphorylated at the 5’ end was ligated to the enriched mRNA. Following the generation of the Illumina TruSeq stranded mRNA library, equal amounts of libraries were pooled and 150 nt single-read sequencing was performed with an Illumina NextSeq 2000 P2 200 cycle kit.

### Public Term-seq datasets

In addition to the *E. coli* Term-seq data, several publicly available datasets were analyzed as part of this work. Datasets originated from various bacterial species, including *E. coli, B. subtilis, L. monocytogenes, E. faecalis, Synechocystis sp*., *S. clavuligerus, S. griseus, S. coelicolor, S. avermitilis, S. lividans, S. tsukubaensis, S. venezuelae*, and *Z. mobilis*. The analyzed Term-seq datasets, together with the description of the samples, reference genome data, and appropriate accession numbers (NCBI Sequence Read Archive) are presented in the Supplementary Table 1. If multiple Term-seq datasets were available for the same species, each dataset has been assigned an abbreviation (a, b, c, d; e.g. *Escherichia coli (a), Escherichia coli (b)*). Those abbreviations are described in the Supplementary Table 1 and will be referred to throughout the paper, particularly in tables and data visualizations. To simplify visualizations, only a single representative study per species is shown (applicable to *E. coli* and *B. subtilis*). Complete visualizations are available in the Supplementary Information.

To present the phylogenetic relationships between analyzed genera (*Escherichia, Synechocystis, Zymomonas, Streptomyces, Bacillus, Listeria* and *Enterococcus*) we downloaded the bacterial reference phylogenetic tree from the GTDB database in the Newick file format (Release 207; available at https://gtdb.ecogenomic.org) (43). The phylogenetic tree was processed using the *etetoolkit* version 3.1.2 (44) to extract only the nodes representing the analyzed genera and *iTOL v6* (45) was used for the tree visualization.

### Analysis of Term-seq data obtained with the protocol by Mondal et al

For each experiment, the quality of the sequencing reads was assessed using *FASTQC* version 0.11.9 (34). Illumina adapters were trimmed using Trim Galore! version 0.6.7 (35). The reads representing 3’ RNA ends (referred to as the `Term-seq dataset`) were selected using the sequence of the Term-seq oligonucleotide ligated to the 3’ RNA ends (sequence: TAGCTCATCGACTGGATCTCAGTGTCTCATT) using *cutadapt* version 1.18 (36) with the *--trimmed-only* option. The minimal overlap between the read and the oligonucleotide to assign to this dataset was set to 8 nt (*-O 8*). Reads shorter than 20 nt were discarded (-*m 20*). The resulting reads were mapped to the appropriate reference genome (see Supplementary Table 1) downloaded from either the *Ensembl Bacteria* version 54 (37) or the *NCBI RefSeq* database (38). Mapping was performed using *bowtie* version 1.2.0 and only uniquely mapped reads were analyzed further (*--best, -m 1* options) (39). Alignments were converted from the SAM file format to BAM, sorted by genomic coordinates, and indexed using *samtools* version 0.1.17 (40). The *bamCoverage version 3*.*5*.*1* (27) from the deep-tools package (with *-- normalizeUsing CPM, --exactScaling, --outFileFormat bigwig, -- binSize 1, --Offset 1* options) was used to generate BIGWIG files representing the normalized genomic coverage of 5’ read ends (corresponding to the 3’ RNA termini). Genomic coverage was calculated separately for each biological replicate and each strand. The signal was normalized by calculating counts per million mapped reads (CPM) values for each genomic position.

The second set, which contains all sequencing reads, was used as the standard RNA-seq dataset. It was processed with *cutadapt* to trim the Term-seq oligonucleotide without discarding reads without a detected overlap. Sequencing reads were mapped to the reference genome using bowtie and the resulting SAM files were processed using *samtools*, as in the case of the Term-seq dataset. Similarly, *bamCoverage* was used to calculate genomic coverage by whole RNA-seq reads, as compared to the Term-seq dataset (excluding the *--Offset 1* option).

*TERMITe find_stable_rna_ends* was run to identify stable 3’ RNA ends separately for each strand using BIGWIG files with the genomic coverage for 3’ RNA ends (Term-seq) and whole RNA-seq reads (RNA-seq) for each replicate using the *--rna-3prime-ends* and *--rna-seq-coverage* options, respectively. Option *--min-no-comp 1* was used to specify that the Term-seq signal needs to be reproducible at least between a single pair of replicates (IDR < 0.05). Next, *TERMITe annotate* was employed to annotate the 3’ RNA ends with *--trans-term-hp, --rnafold, --upstream-nt 100* and *--downstream-nt 10* options. The gene annotations and genomic sequences were used according to Supplementary Table 1. Termination efficiency was calculated only for sites where the average upstream genomic coverage by RNA-seq reads (*--min-upstream)* was higher or equal to the median RNA-seq read coverage for each experiment (see Supplementary Table 1 for exact values).

### Analysis of data obtained with the method published by Dar et al

Similarly, to the protocol described above for data generated with the Mondal et al. protocol, the quality of the reads was assessed using FASTQC, and Illumina adapters were removed using *Trim Galore!* The resulting trimmed sequencing reads were mapped to the reference genome using bowtie and post-processed with *samtools* and *bamCoverage. TERMITe* was run as in the protocol described above, except for the *--rna-seq-coverage* and *--min-upstream options*, which do not apply to data obtained with the Dar et al. protocol due to a lack of matching RNA-seq datasets (14).

### Selection of intrinsic terminators

From the set of stable 3’ RNA ends identified and annotated using the *TERMITe* software, we selected sites corresponding to intrinsic transcription termination sites. We report all intrinsic terminators located within 200 nt downstream from the nearest CDS that were confirmed with either TransTermHP or RNAfold protocol with the additional requirement for those declared solely by the RNAfold pipeline to have the hairpin free energy ≤ -8 kcal/mol.

### Structural analysis of intrinsic terminator hairpins

We analyzed sequences previously identified by the TransTermHP pipeline as terminator hairpins, including potential upstream A-rich and downstream U-rich regions, to generate consensus secondary structures. Hairpin regions from each analyzed experiment were folded by LocARNA (41) and then represented with R2R software (42). Since it is not known whether residues in the U tract basepairs with upstream A/G residues during transcription termination, we removed the lines representing the proposed base pairs from the graphics.

### Statistical assessment of differences in distributions of selected intrinsic terminator features

We compared several features of intrinsic terminators calculated by the *TERMITe* software and their distributions in different bacterial species and lineages. Those features include the hairpin’s Minimum Free Energy (MFE) values, their lengths, scores, and lengths of the hairpin loops, as assigned by the TransTermHP pipeline. The hairpin score can be loosely interpreted as a heuristic approximation of the MFE value (see (32) for the detailed algorithm). The hairpin length was defined as the sum of nucleotides classified as a part of the stem, while the loop length is the number of nucleotides assigned to be a part of the hairpin loop. Additionally, we examined the distributions of the tail scores assigned by the TransTermHP. Briefly, 15 nucleotides immediately downstream from the detected hairpin were assigned a score by the TransTermHP, which considers the U residues located closer to the hairpin to be more important. The tail score (*ts*) is computed using the following equation:

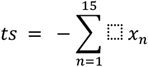

where,

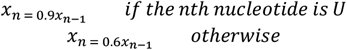

for *n* = 1 …15 and *x*_0_ = 1 (A detailed explanation of both hairpin and tail scores can be found in (32)). Subsequent features include termination efficiencies, distances to the nearest genes located both upstream and downstream on the same strand, as well as the distance between the POT and the base of the predicted terminator hairpin.

We visually examined distributions of measured features either as violin plots or box plots (violin plots being a combination of box plots and kernel density estimates of the underlying distributions) using seaborn version 0.13 (Waskom, 2021). Boxes show quartiles of the distributions, while whiskers extend to 1.5 of the interquartile range. The median is represented with the white dot inside the box. Other visualizations have been prepared using matplotlib (Hunter, 2007) version 3.8 and plotly (Plotly Technologies Inc., 2015) version 5.18.0 libraries. Additionally, we performed a Kruskal-Wallis nonparametric test (function *kruskal* from the scipy.stats Python’s module; scipy version 1.9.2 (28)) to assess whether the samples originated from the same distribution. If the null hypothesis has been rejected at the significance level of p<0.05, the two-sided Mann-Whitney U test (function *mannwhitneyu* from the scipy.stats module; scipy version 1.9.2;) was used to compare the underlying feature’s distributions for each possible pair of samples. Obtained p-values were corrected for the false discovery rate using the Benjamini/Hochberg method (function *fdrcorrection* from the statsmodels.stats.multitest module; statsmodels (46) version 0.13.2). All of the above steps were performed using Python programming language version 3.8.13.

## RESULTS

### *TERMITe* provides robust identification of 3’ RNA ends

*TERMITe* builds on the algorithm implemented in the *termseq-peaks* (26), providing significant modifications in the peak calling procedure and a new module dedicated to annotating the identified stable 3’ RNA ends. Similarly to *Term-seq peaks, TERMITe* software calls peaks in normalized genomic coverage by 5’ Term-seq read ends (Figure 2). Next, both algorithms assign IDR values to the peaks that overlap in pairwise comparisons of replicates. The major difference between tools is that in the next step *termseq-peaks* calculates the number of peaks below the specified IDR threshold for each pairwise comparison and selects a minimal number of such peaks called *N*. The algorithm then merges the signals from all the replicates together, repeats the call for the peaks, and selects the *N* peaks with the highest prominence (Figure 2A). On the contrary, the *TERMITe* software returns all peaks that are found to be reproducible (by default at the significance level of IDR < 0.05) in at least *M* pairwise comparisons of replicates (where the user can specify the parameter M, Figure 2B). Importantly, the selection of the most probable 3’ RNA ends based on prominence in merged replicates (as implemented in *termseq-peaks*) could be strongly biased towards signals from the most highly expressed genes (e.g., rRNA, tRNA, etc.). *TERMITe*, by operating on reproducibility among individual replicate samples, avoids possible biases introduced at this step.

**Figure 2.**
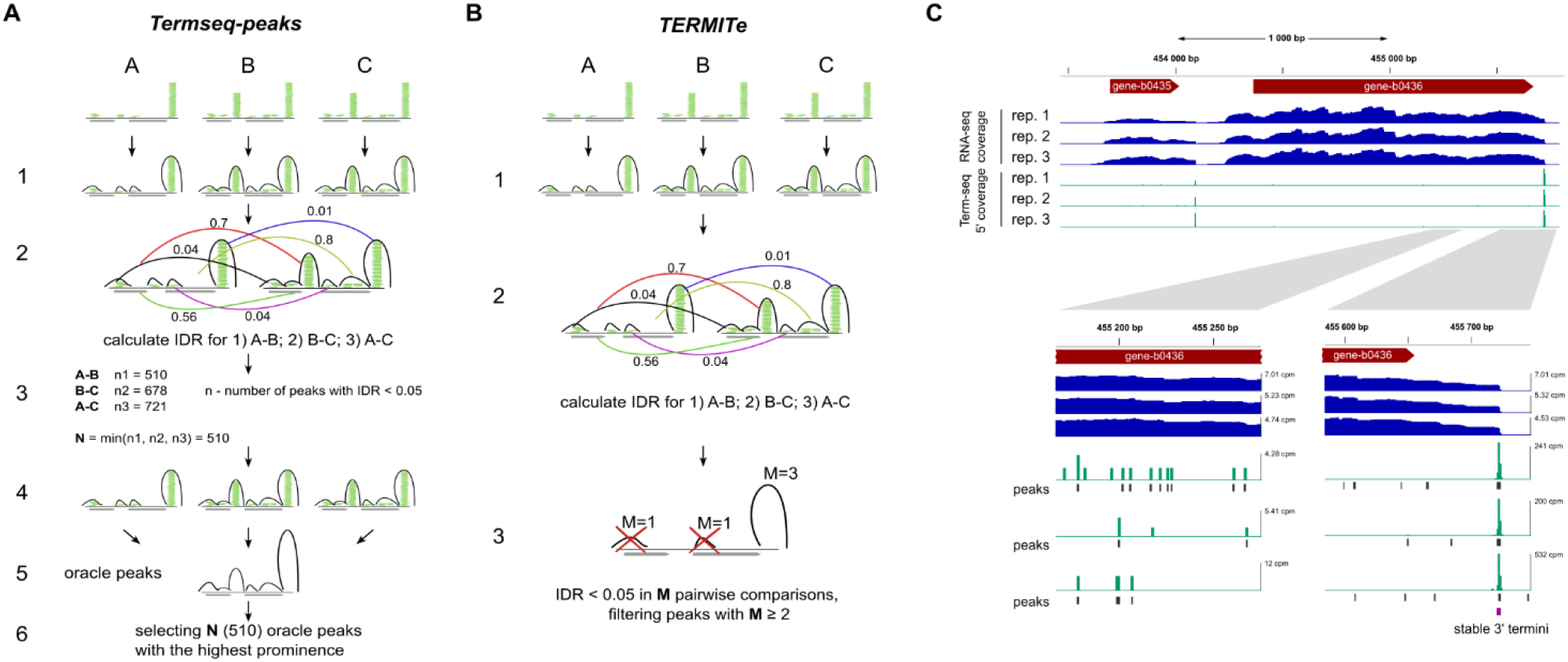
The TERMITe algorithm. **A)** Comparison of the major processing steps included in the *termseq-peaks* and the *TERMITe* methods. *termseq-peaks* pipeline: peaks are called independently for each replicate (A, B, C) with lenient parameters and the prominence is calculated for each peak; IDR is calculated on the basis of prominence values for overlapping peak pairs in each pairwise comparison of replicates; the minimum number of pairs of peaks below the specified threshold (0.05 by default) across all possible pairwise comparisons of replicates is calculated (N); the signal of all replicates is merged and the peaks are called in the combined signal (‘oracle peaks’); N oracle peaks with the highest prominence are selected as final Term-seq peaks. *TERMITe* pipeline: peaks are called independently for each replicate with lenient settings and peak span is trimmed, the prominence is calculated for each peak; IDR is calculated on the prominence values for each pair of overlapping peaks in each pairwise comparison of replicates; peaks with IDR below a specified threshold (0.05 by default) in at least M (user-defined, 1 by default) pairwise comparisons are reported as stable 3’ RNA ends. **B)** Example visualization of the *E. coli* Term-seq and RNA-seq input data, along with *TERMITe* results. The reproducible 3’ RNA termini identified by *TERMITe* have been marked with purple in the bottom right panel. Examples of irreproducible peaks with low and highly varying coverage are presented on the bottom left panel.

In *TERMITe*, the identification of the observed RNA termini (POTs) is followed by downstream annotation, enabling a comprehensive analysis of their biological context. The *TERMITe annotate* module not only provides identification of the genes and genomic features associated with each POT, but also implements a straightforward solution for merging results from multiple samples into a single report. During merging, POTs that overlap between samples are identified and reference positions are estimated based on POT prominence (see the Methods section for details). *TERMITe* also offers a combination of two computational approaches to predict whether a given transcription termination event is related to intrinsic termination. This two-step verification is based on the *TransTermHP* method for predicting intrinsic terminators. However, *TERMITe* significantly increases its sensitivity by implementing secondary structure verification and free energy calculation on a broad set of candidate terminators initially reported using highly relaxed *TransTermHP* search criteria. This approach was feasible due to the limitation of terminator search space to relatively short sequence regions upstream of experimentally identified POTs.

To verify the performance of TERMITe, we decided to analyze the Term-seq datasets from *E. coli*, an organism with the most comprehensive annotation of experimentally verified transcription termination sites. Since the only publicly available *E. coli* Term-seq dataset (17) was prepared with the Dar et al. protocol, we carried out a Term-seq experiment in *E. coli* using the protocol from Mondal et al., to enable a direct comparison of the *TERMITe* efficiency on Term-seq data derived from different experimental protocols. To focus on intrinsic transcription termination events, we only included POTs containing their canonical features and annotated by *TERMITe annotate* as intrinsic terminators. As a result, we identified 691 and 957 intrinsic terminators in the Term-seq data published in this paper and previously published data (17), respectively. The observed difference in the number of identified terminators was primarily due to higher read coverage of the transcript ends in the case of the 3’ end-focused Dar et al. protocol. All the characteristics of the identified terminators (Supplementary Figures 3-11) were highly similar, establishing that *TERMITe* is suitable for the analysis of data derived from both protocols. Thus, *TERMITe* enables users to directly compare stable RNA ends identified for different samples and understand their biological origin, making this the first bioinformatic toolkit that allows a comprehensive analysis of the Term-seq data derived from both experimental protocols.

Next, we evaluated the 3’ RNA ends identified with *TERMITe* against RegulonDb v12.0, a comprehensive database of transcription regulation in E. coli K-12, containing a well-curated set of intrinsic terminators and transcription start sites (TSS) (47). Since the information stored in the database does not represent a complete set of TSS, it was not possible to estimate the software performance measures (specificity, sensitivity) and directly compare the efficiency of *TERMITe* and *termseq-peaks*. We downloaded a set of 349 intrinsic terminators from RegulonDb and checked for overlaps with the set of intrinsic terminators and other stable 3’ RNA ends identified by *TERMITe* in *E. coli*. To perform this task, we added 10 nt flanking regions to all RegulonDb entries and looked for overlaps on the same strand. We observed that 259 out of 349 RegulonDb terminators (∼74%) were also found in the stable 3’ RNA end set of *TERMITe*, specifically 246 in Choe et al. (17) and 219 in the dataset published with this article. Importantly, *TERMITe* annotated 183 of the intersecting 3’ RNA ends as intrinsic terminators. The analysis of distances between intrinsic termination sites found by *TERMITe* to the closest upstream stop codons and the closest downstream transcription start sites from RegulonDB revealed that the majority of termination signals are located within a range of 25 to 75 nucleotides downstream of the nearest stop codon and up to 200 nucleotides upstream from the TSS (Supplementary Figure 1). The observed distances are in an expected range, establishing that *TERMITe* can efficiently annotate intrinsic terminators.

### A comprehensive atlas of intrinsic transcription terminators

Next, we used *TERMITe* to perform a comprehensive analysis of the reported here and publicly available Term-seq datasets to investigate the variability in intrinsic termination signals across bacterial species. In our analysis we included 691 and 957 intrinsic termination sites predicted by *TERMITe* in *E. coli* based on the data described in this study and Choe et al. (17), respectively; 635, 1165, 984 and 1214 sites of *B. subtilis* based on data from Chhabra et al. (15), Mandell et al. (7), Dar et al. (14) and Mondal et al. (8), respectively; 796 terminators from *E. faecalis* (14); 862 terminators from *L. monocytogenes* (14); 206 terminators from *Z. mobilis* (24); 165 terminators from *Synechocystis sp*. (16); 629 terminators from *S. coelicolor* (20), 573 terminators from *S. venezuelae* (20); 520 terminators from *S. tsukubensiis* (20); 737 terminators from *S. griseus* (22); 495 terminators from *S. lividans* (20); 435 terminators from *S. clavuligerus* (21); 707 terminators from *S. avermitilis* (20) (Supplementary Figure 3A). All of the above-listed terminators have been identified using *TERMITe* as described in the Methods section. A complete list of the intrinsic terminators identified in this study is available in Supplementary Table 2. The species analyzed represented relatively high variability in the phylogenetic relationship and GC content (Supplementary Figure 2).

Interestingly, the highest percentage of intrinsic terminators, among all stable 3’ RNA termini identified, was identified in *B. subtilis* (55.6%), followed by *L*. monocytogenes (41.2%) and *E. coli* (37.6%) (Supplementary Figure 3B). This observation indicates that intrinsic termination is a major source of 3’ RNA termini in these species. On the contrary, in *Streptomyces sp*. only a small fraction of POTs met the criteria for canonical intrinsic termination (4-6.6%), suggesting that other pathways for the generation of 3’ RNA termini, such as Rho-dependent termination or RNA processing, could be more prominent in *Streptomyces*. On the other hand, since most of the events were identified solely by the highly sensitive RNAfold pipeline, which is capable of identifying terminators with little or no U tracts (Supplementary Figure 3C), *Streptomyces* might utilize noncanonical intrinsic terminators. To explore those possibilities, we compared numerous features of identified intrinsic terminators.

### Nucleotide composition of intrinsic terminators

First, we analyzed the sequence composition of the intrinsic terminators identified. We observed a significant difference in the nucleotide composition of the U tract region in different bacterial species (Figure 3, Supplementary Figures 4, 5 and 6). The U content in the region after the terminator hairpin in most species was similarly distributed (Figure 3, Supplementary Figures 4 and 5). However, in GC-rich *Streptomyces sp*. it was much lower than in the other bacteria. The number of Us within the first 10 nt downstream of the hairpin in *Streptomyces* sp. was in the range of 3-4, while in most species 7-9 with a peak of 8 Us were observed (Figure 3, Supplementary Figure 6). Only in *E. coli* the number of Us was uniformly distributed in the range of 4-8.

**Figure 3.**
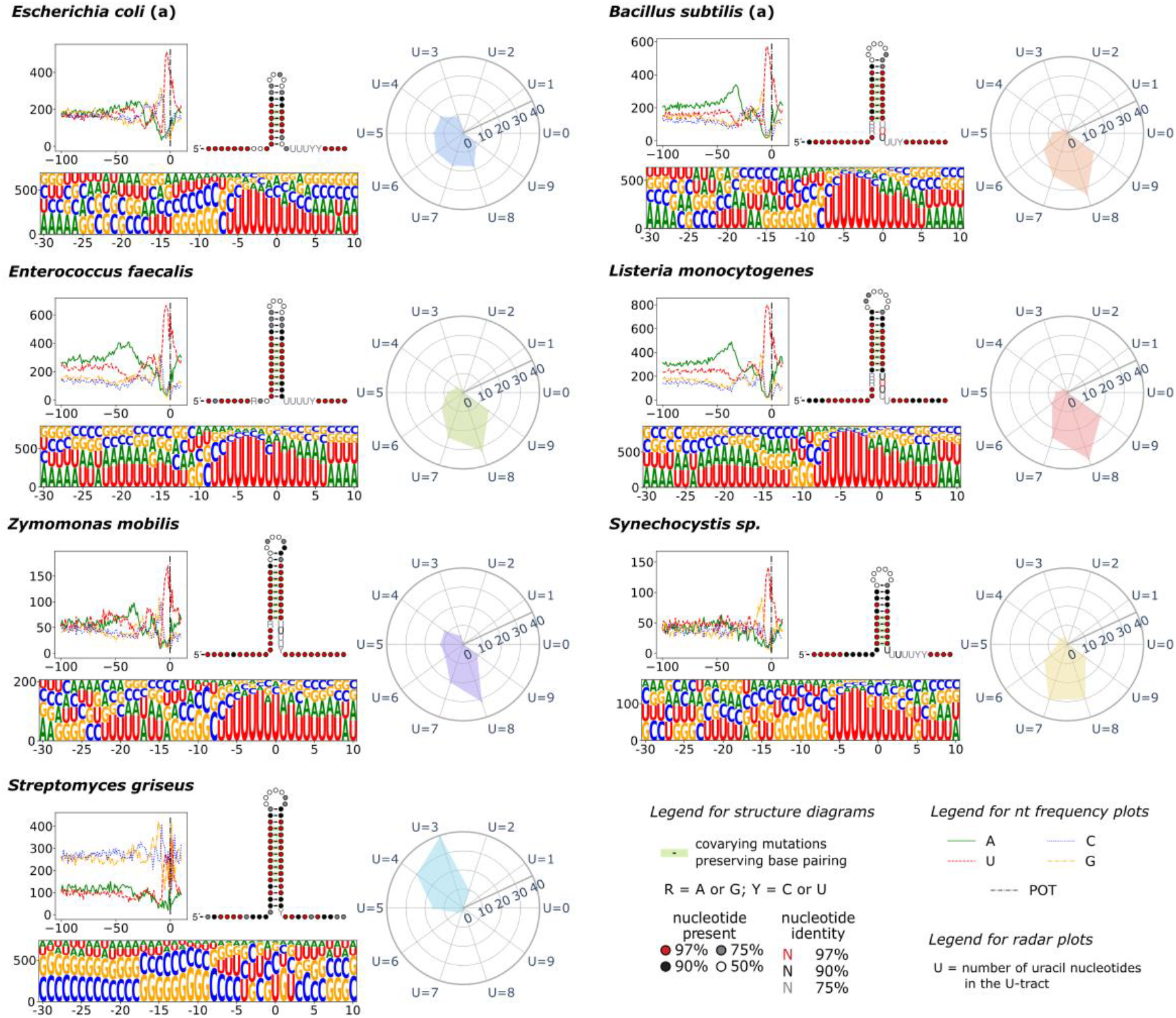
Comparison of distinct features of intrinsic terminators. The top-left plot of each panel shows the nucleotide composition of the sequence regions starting 100 nt upstream and ending 10 nt downstream from the POT for all identified intrinsic terminators in each species. The sequence logo (bottom left of each panel) presents the terminator region (-30 to +10 nt from POT). The height of the letters on the sequence logo is proportional to the frequency of each nucleotide observed at a specific position. The radar plot (right of each panel) shows the percentage of U tracts (defined as ten nucleotides immediately downstream from the hairpin base) that contain the specified number of uracil residues. The consensus secondary structure of the terminator hairpin is also presented, along with information about the conservation of nucleotides and secondary structure elements. The possible base pairing between the U tract and the 5’ A-rich region of the hairpin is removed due to the ambiguous character of such interactions during transcription. The Y-axis on the nucleotide frequency plots and the sequence logo represents the number of terminators with a given nucleotide. For species represented by multiple datasets, only representative results have been shown. For complete results, see Supplementary Figures 4-7.

Similar differences were observed in the distribution of tail scores calculated by the *TransTermHP* pipeline (as described in the Methods section). Differences in tail scores observed between species were found to be significant (Kruskal-Wallis, H-statistic: 2925.97, p-value: <0.001). Not surprisingly, we found that intrinsic terminators in the *Streptomyces* genus were assigned considerably higher tail scores (median range -3.6 to -3.4), while the other bacterial species exhibit significantly lower tail scores (median range -5.5 to -4.7) (Figure 4B, Supplementary Table 8). Also, in accordance with the above analysis of the distribution of Us, we identified considerably more terminators with relatively weak U tracts in *E. coli* (third quartile (Q3) = -3.8; -3.9) as compared to *B. subtilis* (Q3 = -4.7; -4.6; -4.7; -4.6), *L. monocytogenes* (Q3 = -5.1), *Z. mobilis* (Q3 = -4.4) and *Synechocystis* (Q3 = -4.9). The detailed results of all pairwise comparisons of samples using the Mann-Whitney U test are shown in Supplementary Table 4. The above observations confirm the noncanonical character of *Streptomyces* U tracts, which could interfere with efficient annotation of identified POTs as intrinsic terminators.

**Figure 4.**
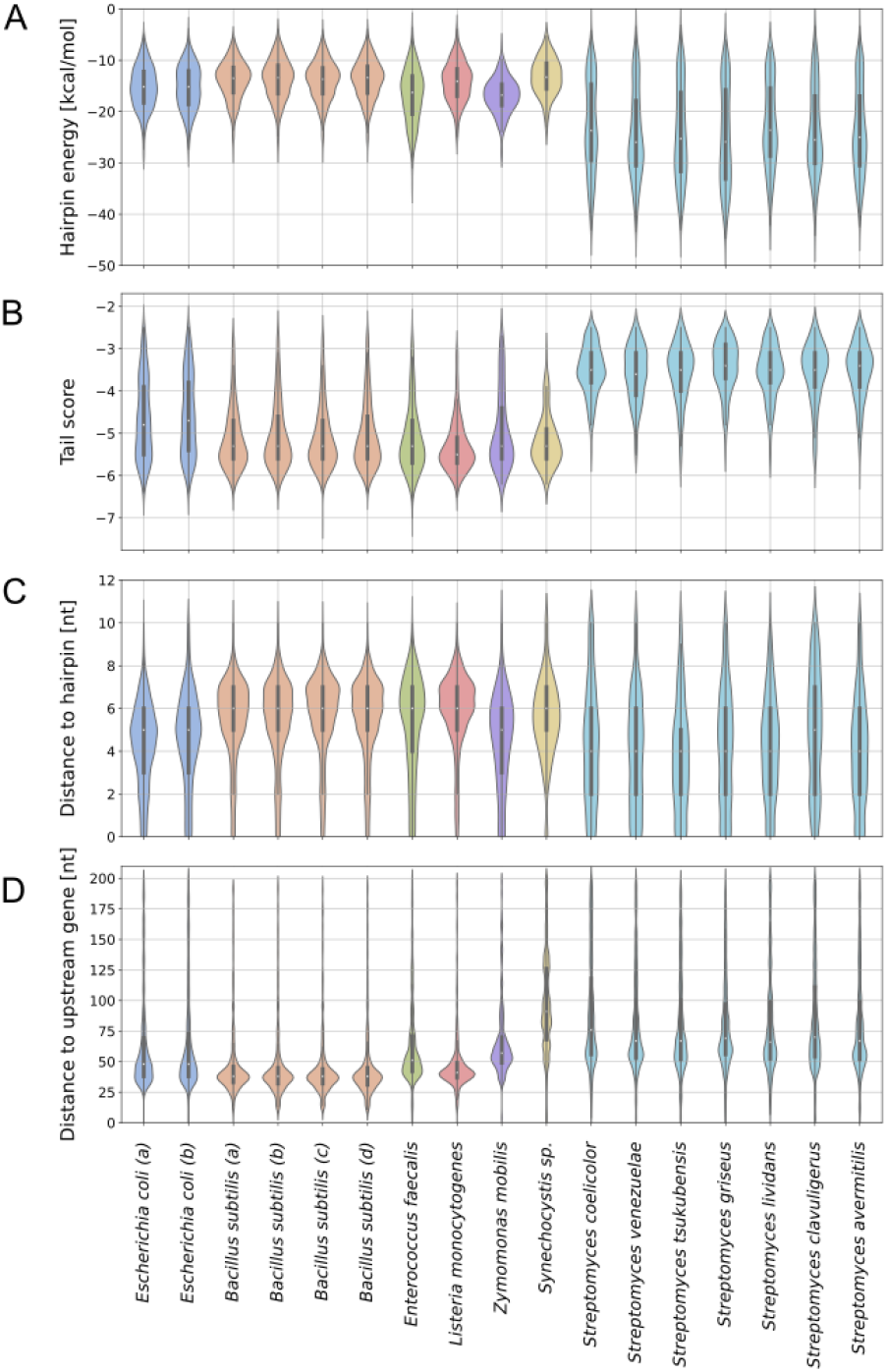
Comparison of the distinct features of intrinsic terminators in various bacterial species. **A)** Distributions of hairpin free energies assigned by *TERMITe* to intrinsic terminators. **B)** Distributions of the tail scores assigned by *TERMITe* to intrinsic terminators identified by the TransTermHP pipeline for each experiment and species analyzed. **C)** Distributions of the distances between the hairpin base and the POT for intrinsic terminators identified by *TERMITe*. **D)** Distributions of the distances between the POT and the nearest gene upstream on the same strand.

Interestingly, in *B. subtilis, L. monocytogenes, E. faecalis*, and to a more limited extent in *Z. mobilis* and *E. coli*, we observed significant enrichment in adenines (A-rich tracts) in a region from -25 to -50 upstream of the POT (Figure 3, Supplementary Figure 4). This is an essential feature of bidirectional terminators capable of terminating transcription on both strands of DNA (48). The prevalence of A-rich tracts was much higher in *B. subtilis, E. faecalis* and *L. monocytogenes* than in *E. coli* and virtually absent in *Synechocystis* and *Streptomyces* sp. Additionally, in all species analyzed, we observed a specific increase in GC content within the hairpin region with a specific high peak in GC or G directly before the beginning of the U tract.

### Structural features of terminator hairpins

Next, we compared the structural features of the intrinsic terminators associated with identified POTs. MFE values of identified terminator hairpins tend to have substantially lower free energies (more stable hairpins) in *Streptomyces* sp. (medians range from -25.95 to -23.6 kcal/mol) than in other bacteria (medians range from -16.8 to -13.3 kcal/mol) (Figure 4A, Supplementary Table 3). The observed differences in MFE population medians for different bacteria are statistically significant (Kruskal-Wallis, H-statistic: 2857.70, p-value: <0.001, Supplementary Table 3). Interestingly, we also observed an increase in the number of terminators with relatively low MFE values in *E. faecalis* (first quartile (Q1) = -20.5 kcal/mol), and to some extent in *E. coli* (Q1 = -18.3 kcal/mol; -18.6 kcal/mol) and *Z. mobilis* (Q1 = -18.8), compared to *B. subtilis* (Q1 = -16.3 kcal/mol; -16.5 kcal/mol; -16.5 kcal/mol; -16.4 kcal/mol), *L. monocytogenes* (Q1 = -17.0 kcal/mol) and *Synechocystis* (Q1 = - 15.6 kcal/mol) (Figure 4A, Supplementary Table 3). These differences in the MFE distribution were statistically significant except for comparisons of *E. faecalis* – *Z. mobilis* (FDR = 0.57), *Synechocystis sp*. – *B*.*subtilis* (FDR = 0.1), and *L. monocytogenes* – *B. subtilis* (FDR = 0.07; Mann-Whitney U test with the significance level of FDR < 0.05, Supplementary Table 4) reflecting the above differences. Similar conclusions were drawn during the examination of the TransTermHP hairpin score distributions (Supplementary Figure 8, Supplementary Table 5), which can be interpreted as a heuristic approximation of the MFE values.

We also observed significant differences in hairpin lengths between species (Kruskal-Wallis, H-statistic: 3207.36, p-value: <0.001; Supplementary Figure 9, Supplementary Table 6). Terminator hairpins in *Streptomyces* sp. are considerably longer (medians in the range of 32 to 35 nt) than in other bacterial species (medians range of 20 to 28 nt), except *E. faecalis*, which is characterized by a similar distribution of hairpin lengths (median 33 nt; Supplementary Figure 9, Supplementary Table 6, Supplementary Table 4). However, the hairpin loop lengths were highly consistent with a median of 4 for each species (Supplementary Figure 10, Supplementary Table 7, Supplementary Table 4).

### Localization of termination sites

Next, we have analyzed the positions of the identified intrinsic Next, we analyzed the positions of the POT relative to terminator features. In all species, the POTs were located in the central part of the U tract reaching positions of -6 to +7 in *E. coli*, -7 to +5 in *B. subtilis*, - 8 to +6 in *E. faecalis*, -8 to +7 in *L. monocytogenes*, -7 to +8 in *Z. mobilis*, -6 to +4 in *Synechocystis* sp, and –7 to +6 in *Streptomyces* relative to POT (Figure 3, Supplementary Figures 4 and 5). We also identified three clusters of bacterial species by visually inspecting the distributions of the distances between the POT and the hairpin base. We observed the smallest average distance for the *Streptomyces* genus (medians of 4, except for *S. clavuligerus* with the median of 5), intermediate distances in Gram-negative species (medians of 5 nt for *E. coli* and *Z. mobilis*, and 6 nt for *Synechocystis*), while highest average distances among the bacterial species analyzed were observed in Gram-positive bacteria (medians for *B. subtilis, L. monocytogenes, E. faecalis* = 6; Figure 4C; Supplementary Tables 4 and 11). The differences in distance between the POT and the base of the predicted terminator hairpin were statistically significant (Kruskal-Wallis, H-statistic: 1104.15, p-value: <0.001) (Supplementary Table 11). Importantly, all these distances are shorter than the 7-9 nt spacing observed by in vitro transcription using *E. coli* and *B. subtilis* RNAP (6–8). The observed shorter distances may reflect trimming by 3’ to 5’ exonucleases.

Important differences between species were also observed in the location of the intrinsic terminators relative to the closest upstream CDS, reflecting the lengths of the 3’ UTR regions. In particular, in *Synechocystis* and *Streptomyces*, the transcript ends tend to be further from the closest upstream CDS (medians for Synechocystis 91 nt; median for *Streptomyces* in the range of 66 to 76 nt) than in other bacteria (medians range from 38 to 57 nt; Figure 4D). The smallest median distances were observed in *B. subtilis* and *L. monocytogenes* with median distances of 38 nt and 37 nt, respectively. *E. coli, E. faecalis, and Z. mobilis w*ere characterized by slightly longer distances (medians for *E. coli* of 48 nt, for *E. faecalis* of 42 nt, and *Z. mobilis* of 57 nt, respectively). The differences in distances between the POT and the nearest upstream CDS were statistically significant (Kruskal-Wallis, H-statistic: 3566.80, p-value: <0.001) (Figure 4D, Supplementary Table 9).

We also examined the distances between the POT and the closest downstream genes located on the same strand (Supplementary Figure 11, Supplementary Table 10). Differences in population medians for the investigated species were found to be statistically significant (Kruskal-Wallis, H-statistic: 96.16, p-value: <0.001), although the differences were relatively small compared to the other characteristics analyzed. Detailed results of all pairwise comparisons of samples using the Mann-Whitney U test are available in Supplementary Table 4.

### Termination efficiency of intrinsic terminators

Finally, we compared the termination efficiencies calculated for *E. coli* and *B. subtilis* using samples obtained with the protocol published by Mondal et al. (8) for which the matching RNA-seq data were available (Supplementary Figure 12, Supplementary Table 12). The observed efficiencies were high (median in a range of 83.2% to 96.7%) reflecting the high efficiency of the intrinsic terminators in those species. Although we found that the differences in efficiency distributions are statistically significant (Kruskal-Wallis, H statistic: 301.13, p-value: <0.001), it is not possible to conclude that they reflect important variability between species, since the variability for different datasets in the same species (*B. subtilis*) was higher than between *B. subtilis* and *E. coli* (Supplementary Table 12).

## DISCUSSION

Despite recent advances in developing high-throughput protocols tailored to sequencing and identifying 3’ RNA ends in bacteria, a comprehensive bioinformatics tool dedicated to analyzing such data was missing, hampering advances in research on transcription termination in bacteria. The only available tool, *termseq-peaks* was developed for the analysis of Term-seq data obtained with the protocol by Dar et al., which is characterized by substantial enrichment of the cDNA library in RNA 3’ ends (26). The algorithm presented in this work is a universal method designed and tested on data derived from both available Term-seq protocols developed by Dar et al. and Mondal et al. The *TERMITe* provides important advancements by reducing the expression bias in the identification of POTs, utilizing matched RNA-seq data for calculation of termination efficiency, and providing robust annotation of the identified sites. The ability of *TERMITe* to distinguish intrinsic termination events from other 3’ RNA termini enables its effective employment in research projects focused on transcription termination.

A few databases encompassing computationally predicted intrinsic terminators in bacterial genomes have already been published, including WebGeSTer (49) and TransTermHP (32). However, the amount of experimentally validated terminators remains scarce and is limited only to a few model species. For example, RegulonDB (47) and EcoCyc (50) are examples of databases gathering information on experimentally verified terminators for *E. coli*. Considering the diversity of bacteria and the roles they play in human health and agriculture, it is essential to better understand all aspects of gene expression to improve our ability to rationally engineer bacteria to our advantage. Therefore, the reported in this work atlas of intrinsic terminators identified by analysis of experimental Term-seq datasets, containing thousands of terminators found in 13 distinct bacterial species and lineages, provides a valuable resource to the field. This atlas constitutes the most comprehensive dataset of experimentally verified intrinsic terminators, allowing researchers to better understand the mechanisms of transcription termination in bacteria.

Our analysis of identified intrinsic terminators reveals significant variability in the nucleotide compositions of terminator hairpins, U-rich and A-rich tracts, as well as in the distributions of hairpin free energy values, distances to the upstream CDSs and downstream genes, and the distances between the observed POTs and the corresponding hairpin. The most striking differences were observed in bacteria from *Streptomyces* species, which are characterized by extremely GC-rich genomes. Based on inspection of the U tracts located within 10nt downstream of the stem, the *Streptomyces* genus was characterized by the smallest enrichment in U residues. *E. coli* was observed to have much higher enrichment, while the highest was observed for the rest of the analyzed bacterial species. The rarity of the strong U-rich tracts in *Streptomyces* genomes was noted previously (18–22). It has also been reported that terminators with U-rich tracts tend to terminate more strongly in *S. avermitilis* (18). Interestingly, the observed U tract length and strength seem to be negatively correlated with hairpin energy and length. In *Streptomyces*, which is characterized by the weakest U tracts, the hairpins are the most stable and the longest. Long hairpins suggest that their extraordinary stability should not be considered as a side effect of high GC content in *Streptomyces* genomes, but rather an effect of evolutionary pressure. In *E. coli*, which contains a wider distribution of U tract lengths also has a wider distribution of hairpin energy compared to e.g. *B. subtilis*. Such dependency raises questions about possible mechanisms of compensation of weak U tracts by more stable and longer hairpins.

In contrast to the other analyzed bacteria, the free energy values of the intrinsic terminator hairpins in *Streptomyces* sp. follow a bimodal distribution. It was previously suggested that such distribution is related to the evolution of two distinct mechanisms of transcription termination (Rho-dependent and Rho-independent/intrinsic) (16, 21, 22). Thus, the observed bimodality might suggest that the Rho-dependent mechanism might be utilized more often in the GC-rich *Streptomyces* genus than in other bacteria, having less GC-rich genomes, which is in agreement with the observed lowest fraction of intrinsic terminators among all identified 3’ RNA termini. Additionally, in *Streptomyces* the POTs tend to be located closer to the base of the stem than in other bacterial counterparts, suggesting that there are noticeable differences in the mechanisms of transcription termination and/or posttranscriptional RNA processing. Intrinsic terminators in the *Streptomyces* genus were also found to be located further downstream from the 3’ ends of the CDS than in other lineages, except for *Synechocystis*. However, this distinction might not be representative because of the relatively small number of identified intrinsic terminators.

The analysis of the nucleotide composition of terminator regions revealed clear A-rich tracts located immediately upstream from the terminator hairpin stems (25-50 nt upstream of the POTs). It has come to our attention that apart from their absence in *Synechocystis* and *Streptomyces*, A-rich tracts are far less prevalent in the Gram-negative bacteria than in their Gram-positive counterparts. Such tracts are thought to be a characteristic feature of bidirectional transcription terminators (48). It was also suggested that the upstream A-tracts may contribute to the stability of unidirectional terminator hairpins by additional pairing with the U tracts (51). Another possibility could be that the upstream A tracts may serve as structural insulators not allowing the formation of the local secondary structure competing with the folding of the termination hairpin. However, their preferential occurrence in Gram-positive bacteria observed in our analysis remains elusive.

To our knowledge, this is the first multi-species comparison of intrinsic terminator features based on experimentally determined transcription termination sites. In contrast to the previous analyses based on computational predictions, the involvement of Term-seq data allowed us to include non-canonical termination signals, providing an unbiased dataset of transcription termination sites. Our results will facilitate a better understanding of the mechanisms underlying intrinsic termination and underscore the observation that their features might not be as well conserved within the bacterial domain as previously thought. The *TERMITe* software presented in this work provides an important bioinformatic tool to facilitate research on transcription termination using high-throughput datasets obtained with Term-seq.

## AVAILABILITY

*TERMITe* was implemented in Python 3 and containerized with Docker. The software can be downloaded freely from the GitHub repository (*https://github.com/zywicki-lab/TERMITe*).

## ACCESSION NUMBERS

Sequencing data are publicly available in the NCBI Sequence Read Archive with the accession number PRJNA906280.

## SUPPLEMENTARY DATA

Supplementary Data are available online.

## AUTHOR CONTRIBUTION

JGK and MŻ designed the TERMITe algorithm; JGK implemented the algorithm and performed computational analyses; SR and PB created E. coli Term-seq libraries; AChP performed the analysis of secondary structure conservation; AChP, JGK and MŻ prepared data visualizations; JGK, MŻ, PB and MK interpreted the results and evaluated the algorithm; JGK and SR wrote the initial draft of the manuscript; MŻ, MK, PB and JGK wrote the final version of the manuscript.

