## Supplementary Figures and Tables for "Characterization of bacterial intrinsic transcription terminators identified with TERMITe – a novel method for comprehensive analysis of Term-seq data"

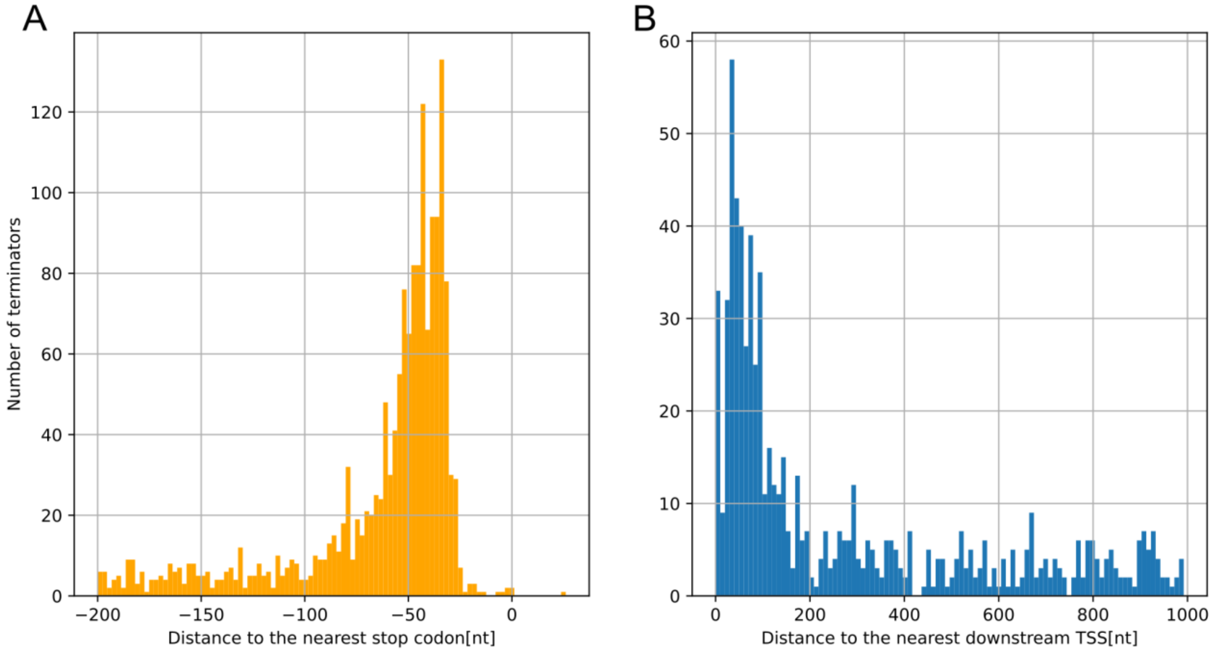


**Supplementary Figure 1.** Distribution of distances from identified intrinsic termination sites to closest genes. **A)** Distances to closest upstream stop codon. **B)** Distances to closest downstream transcription start site.

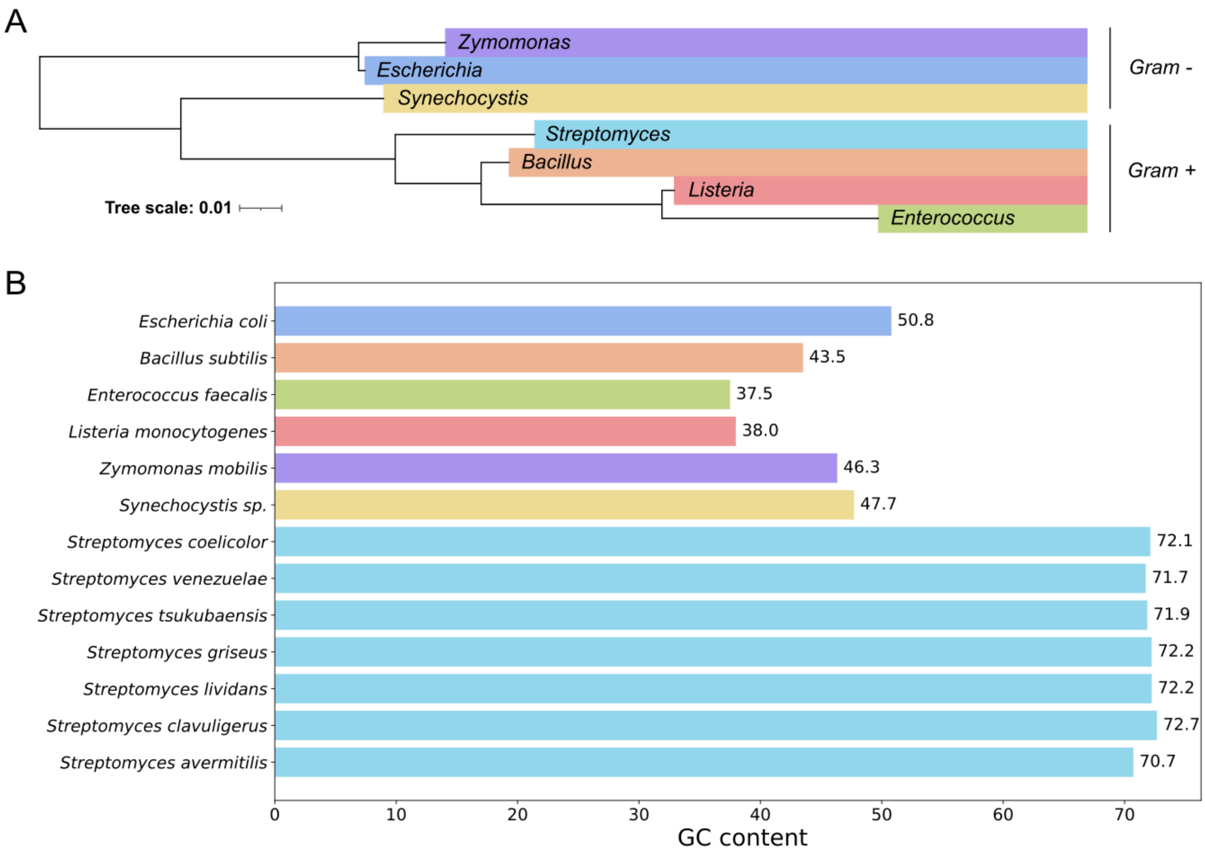


**Supplementary Figure 2. A)** Phylogenetic tree of the analyzed genera. Gram-positive bacteria are marked with (+), while Gram-negative with (-). The phylogenetic tree was visualized using *iTOL v6* (Letunic and Bork, 2021). See Materials and Methods for detailed description of the visualization procedure. The lengths of the edges correspond to the evolutionary distances between genera**. B)** %GC content of the analyzed reference genomes (plasmid sequences were not considered).

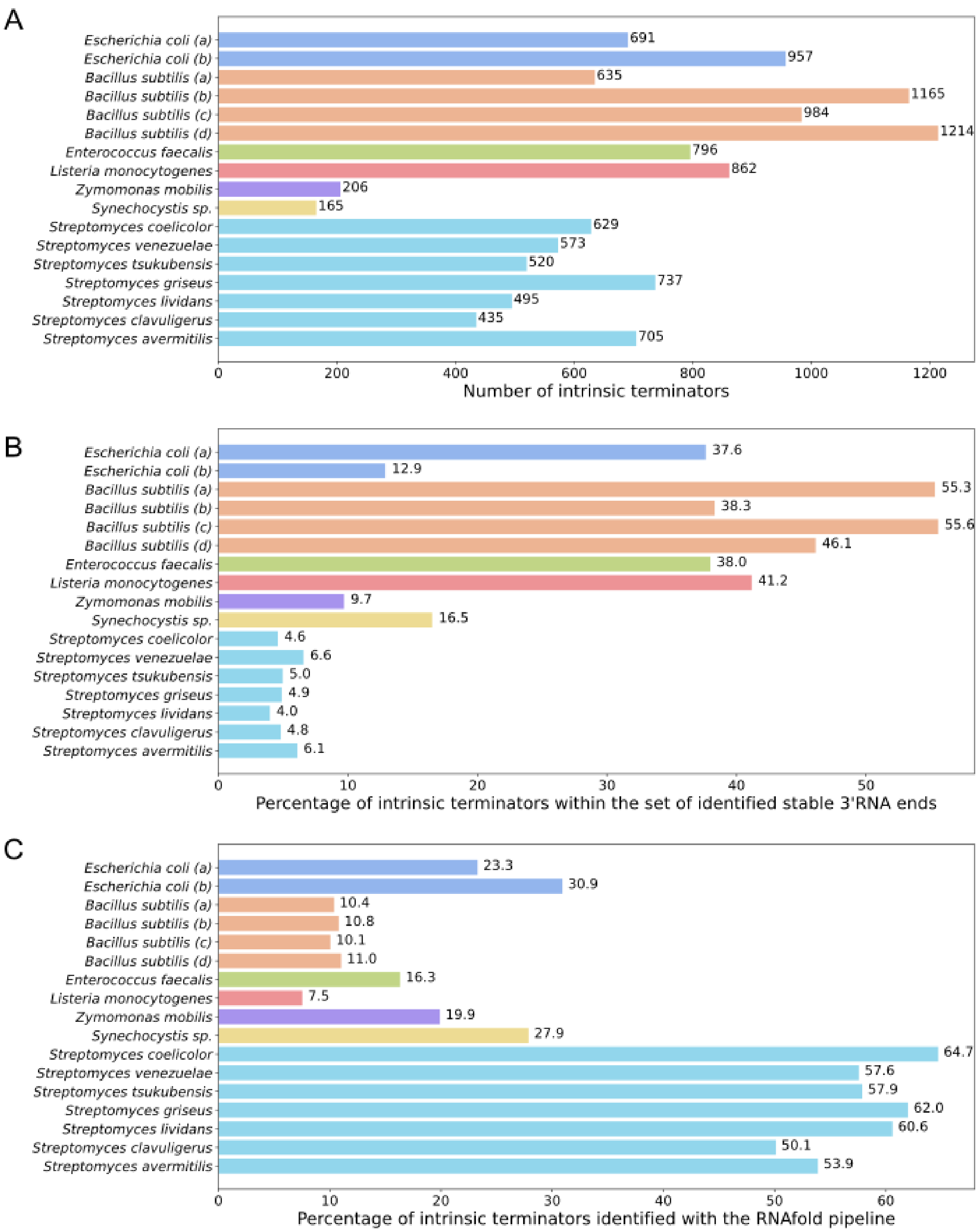


**Supplementary Figure 3.** Summary of intrinsic termination events identified by TERMITe**. A)** The number of identified intrinsic transcription terminators. **B)** The percentage of the stable 3’ ends reported as intrinsic terminators for each species and experiment. **C)** The percentage of intrinsic terminators identified solely with the RNAfold pipeline.

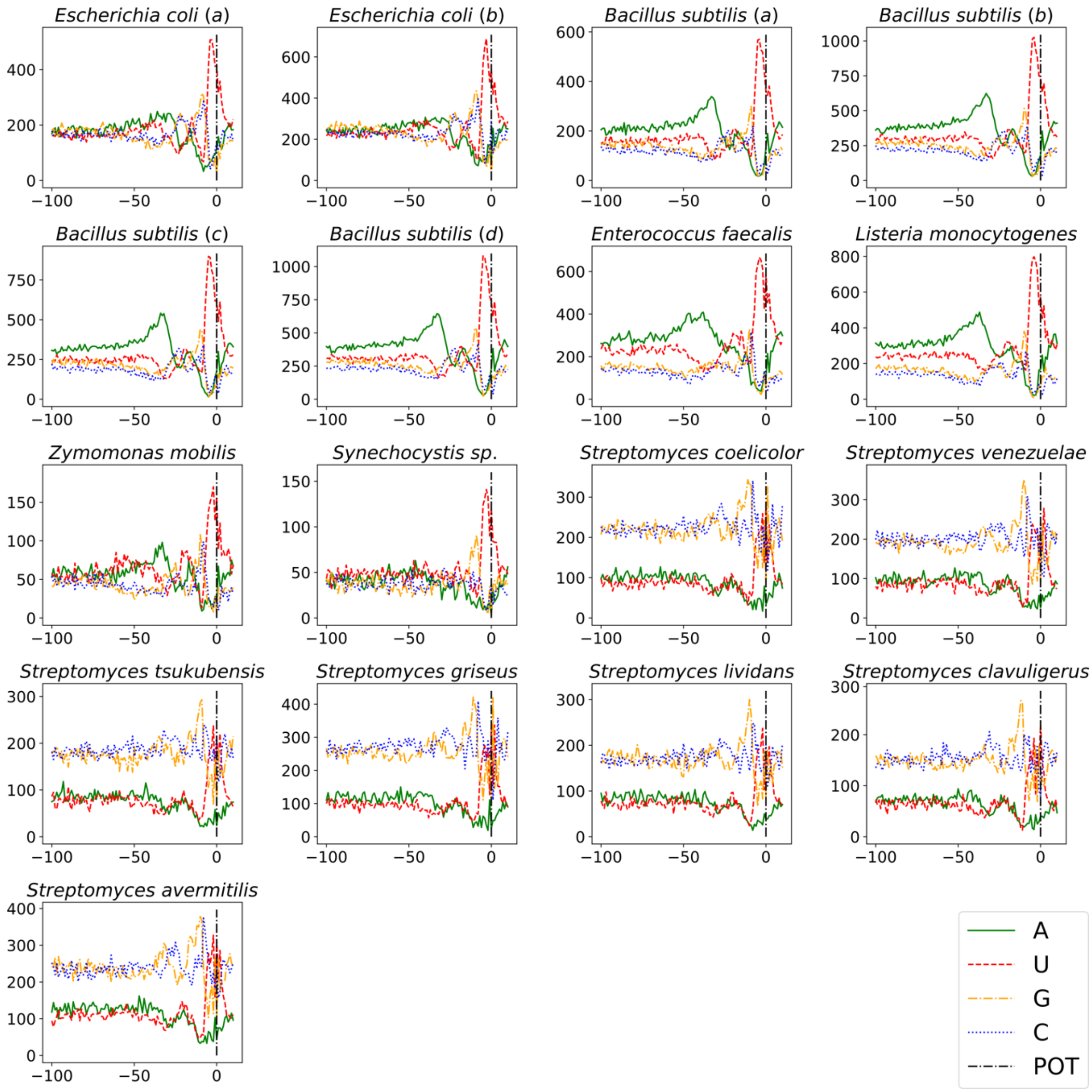


**Supplementary Figure 4.** Nucleotide composition of the sequence region starting 100 nt upstream and ending 10 nt downstream from the POT (position 0) for all identified intrinsic terminators in each experiment. The black vertical line marks the POT.

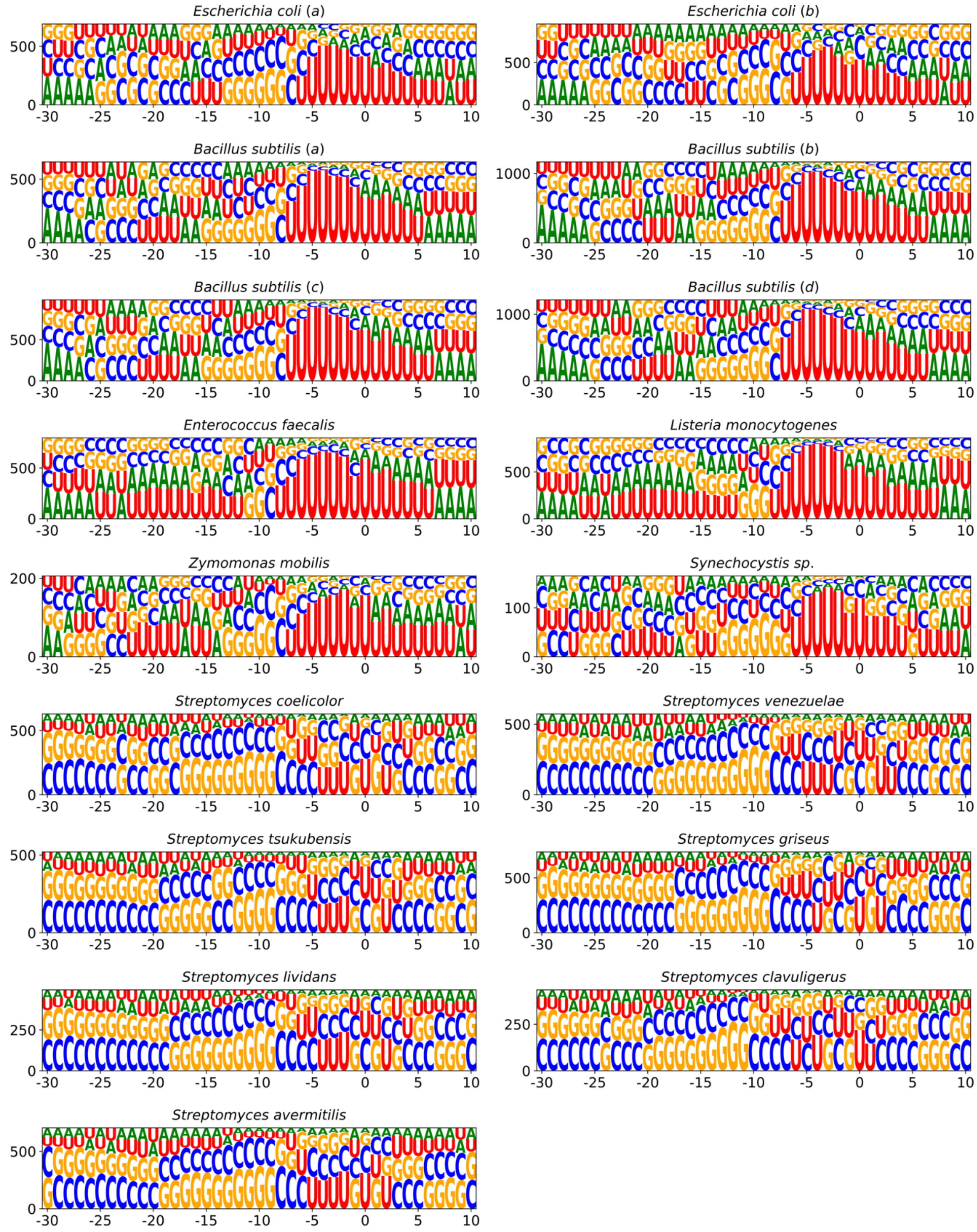


**Supplementary Figure 5.** Sequence logos showing the nucleotide compositions of the terminator regions (30 nt upstream and 10 nt downstream from the POT) for all identified intrinsic terminators in each experiment. The POT is at position 0.

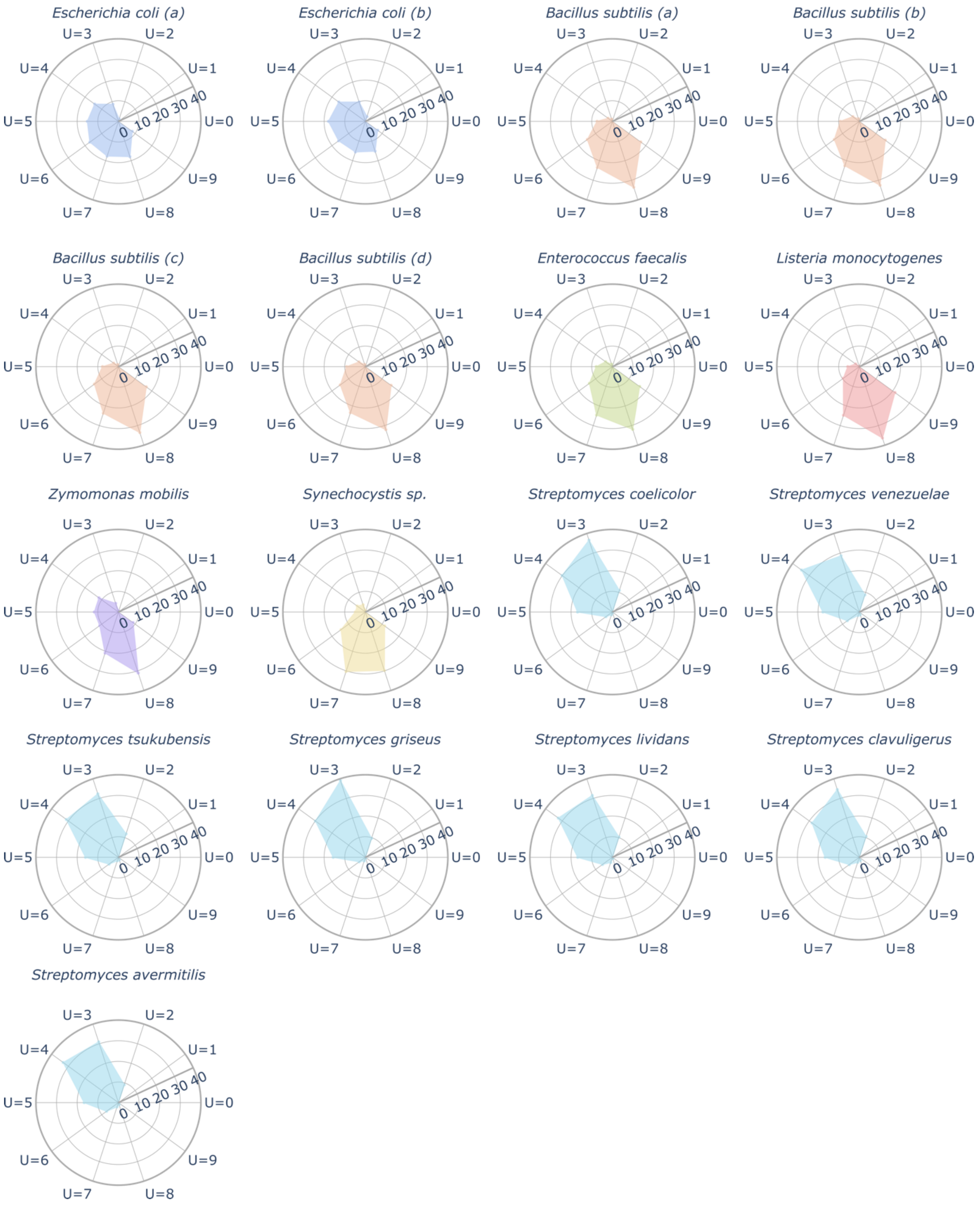


**Supplementary Figure 6.** The percentage of the U tracts (defined as 10 nucleotides immediately downstream from the hairpin base) containing the specified number of U nucleotides within all identified intrinsic terminators in each experiment.

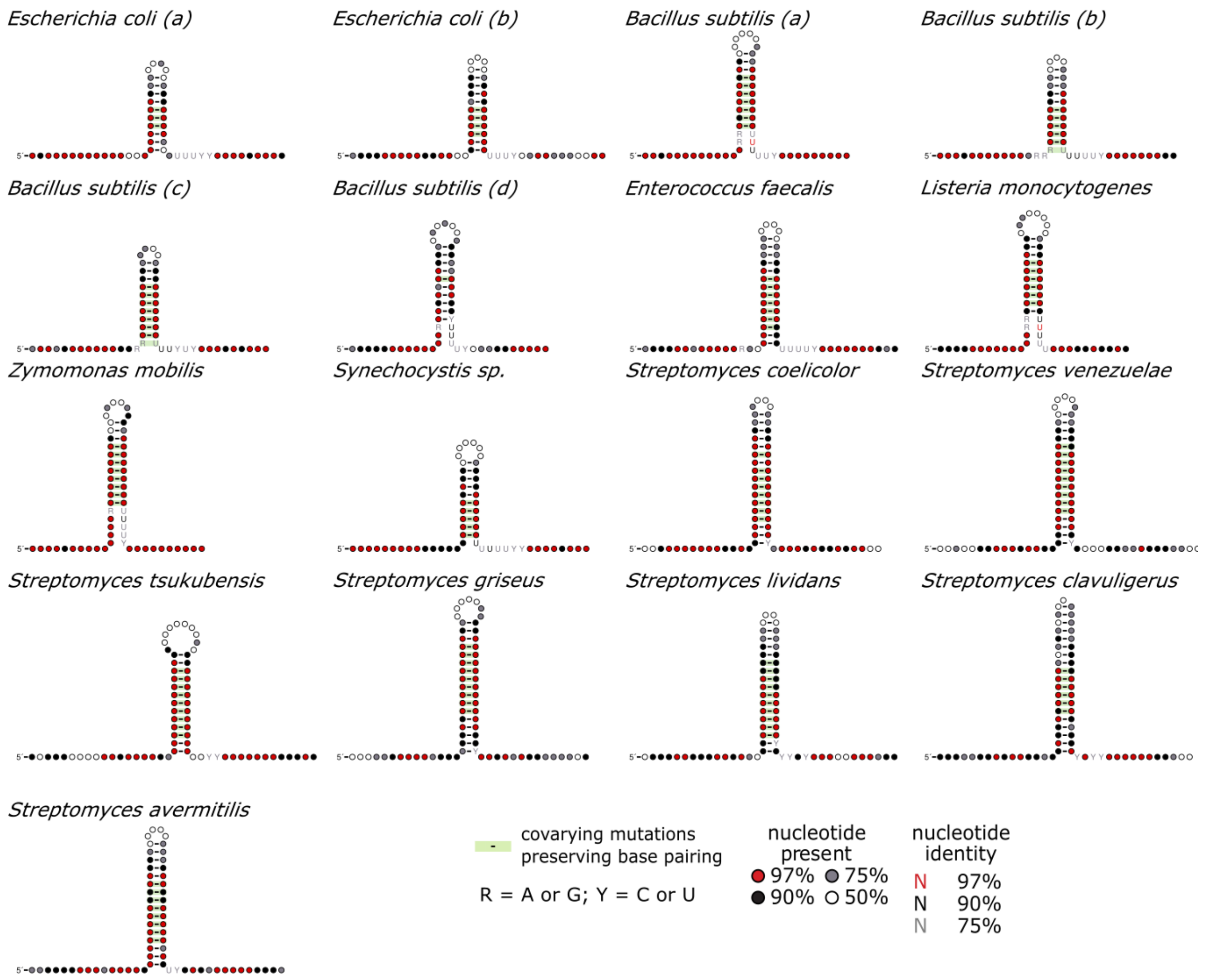


**Supplementary Figure 7.** The consensus secondary structure of all identified intrinsic terminator hairpins along with the information about nucleotide conservation. The base-pairing between the U-rich tract and the 5’ region of the hairpin was unmarked manually. See detailed description can be found in Figure 3 in the main text.

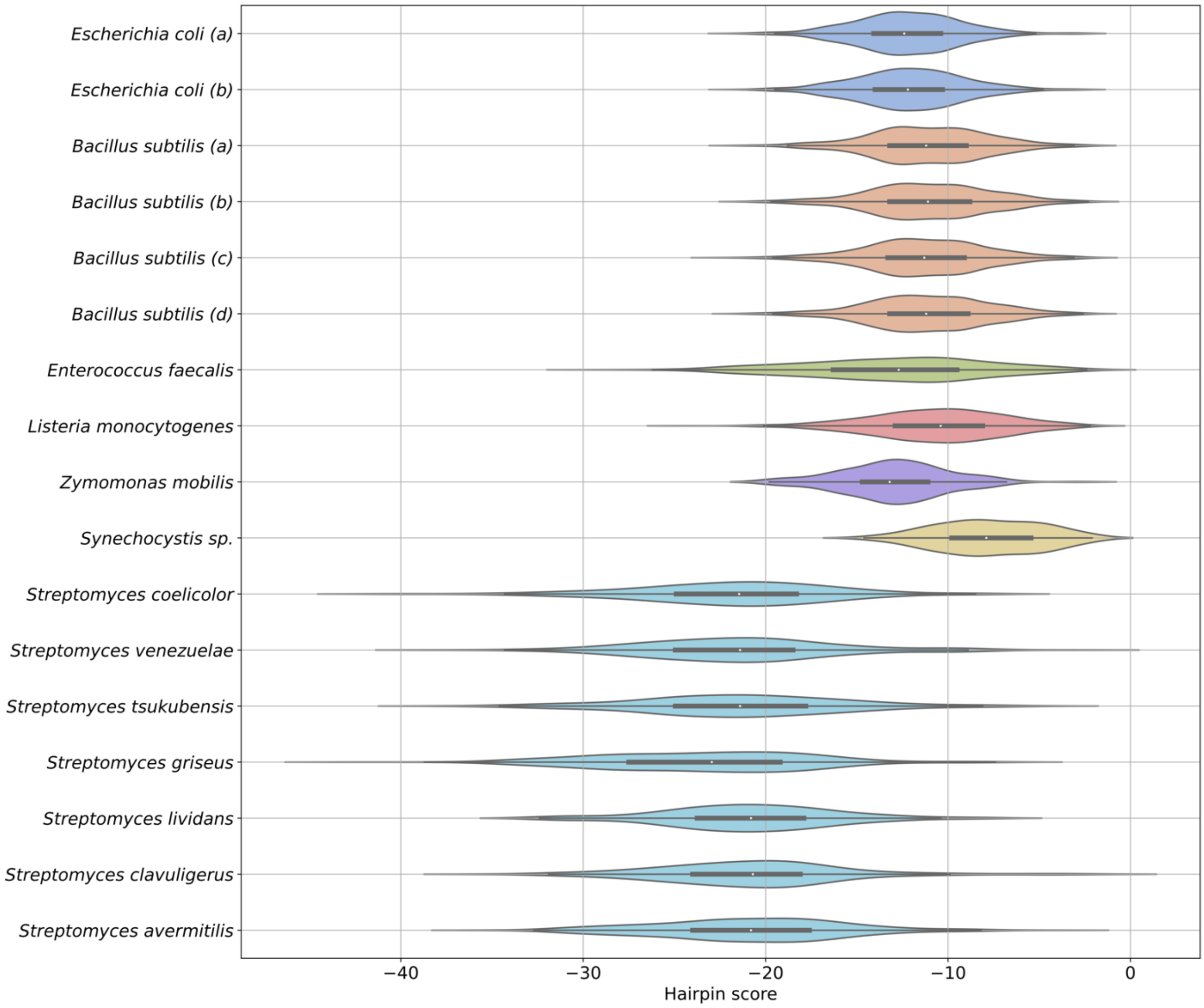


**Supplementary Figure 8.** Distribution of hairpin scores calculated by the TransTermHP pipeline for all identified intrinsic terminators in each dataset.

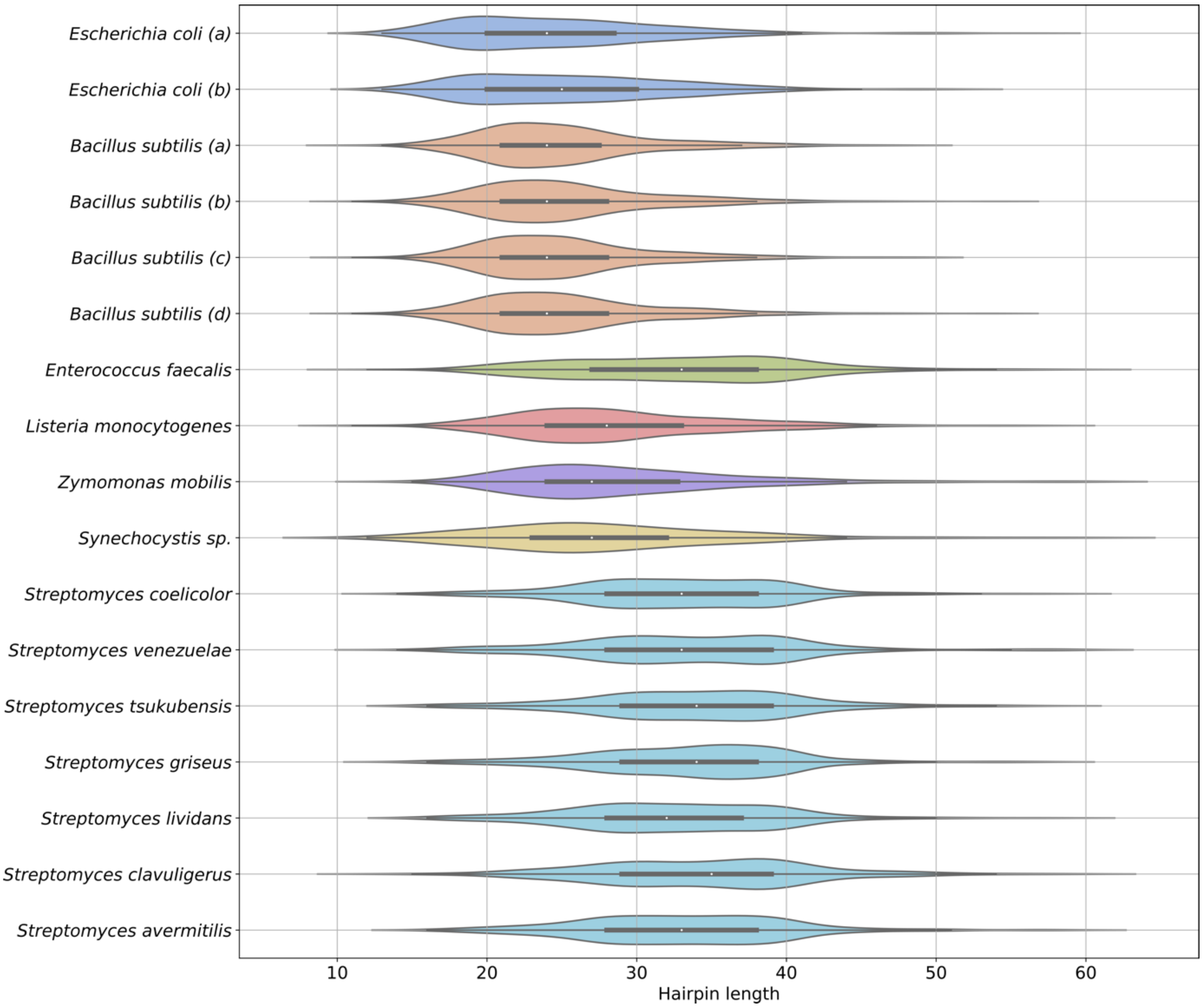


**Supplementary Figure 9.** Distribution of hairpin lengths of all identified intrinsic terminators in each experiment. The hairpin length is defined as the number of nucleotides involved in the hairpin formation.

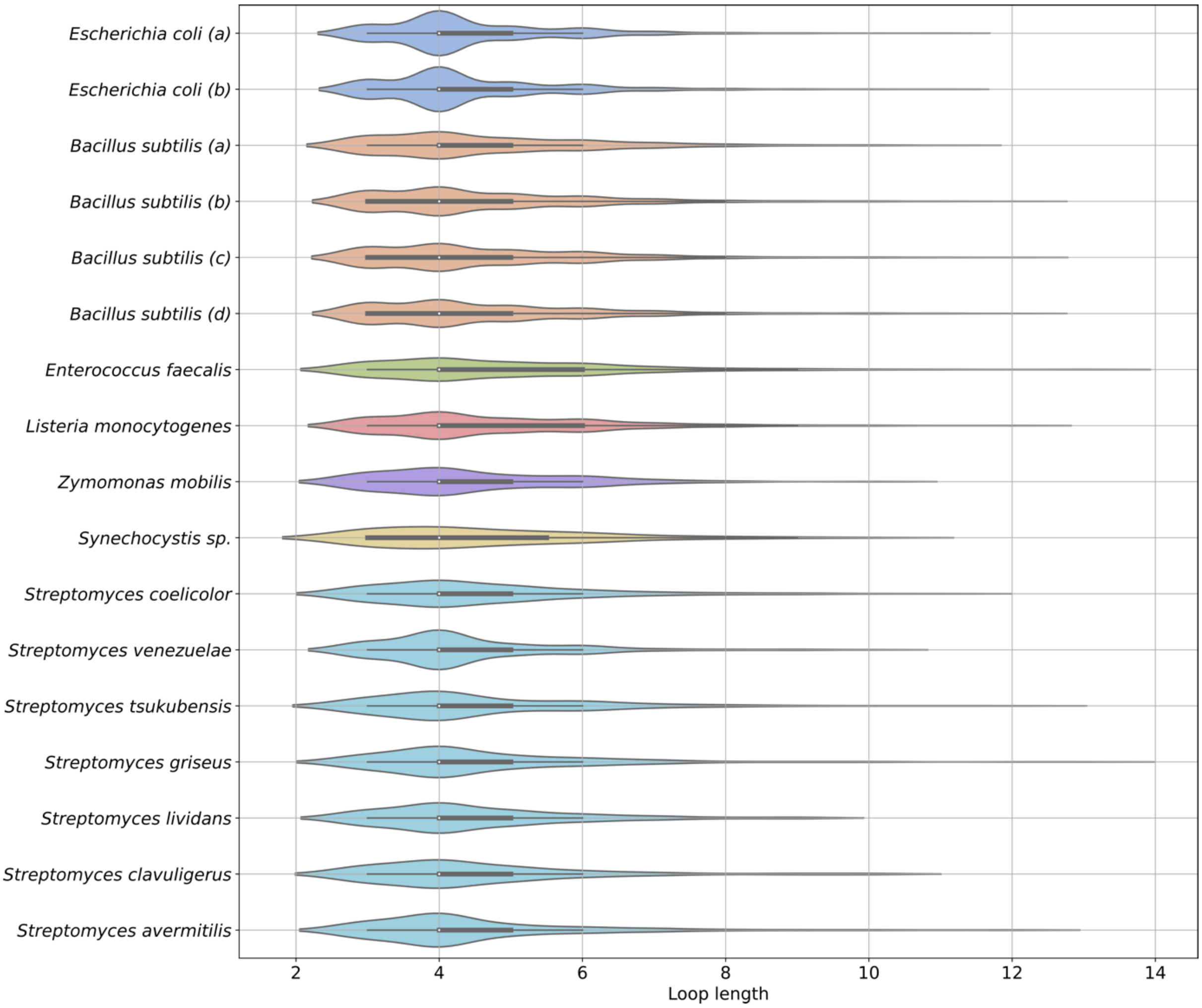


**Supplementary Figure 10.** Distribution of hairpin loop lengths of all identified intrinsic terminators in each analyzed dataset.

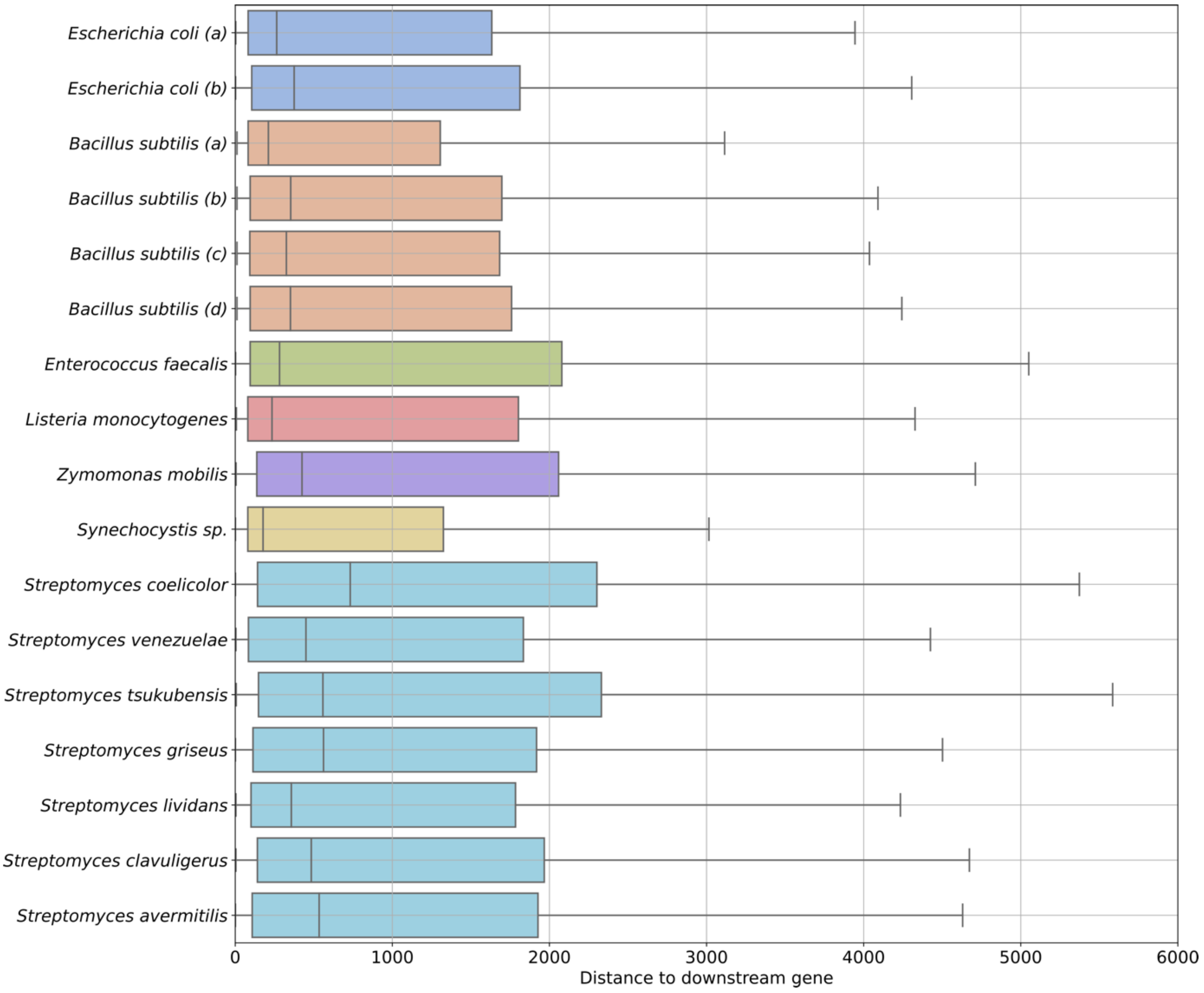


**Supplementary Figure 11.** Distributions of the distances between the POT and the nearest downstream CDS located on the same strand. Distributions are presented separately for each analyzed dataset.

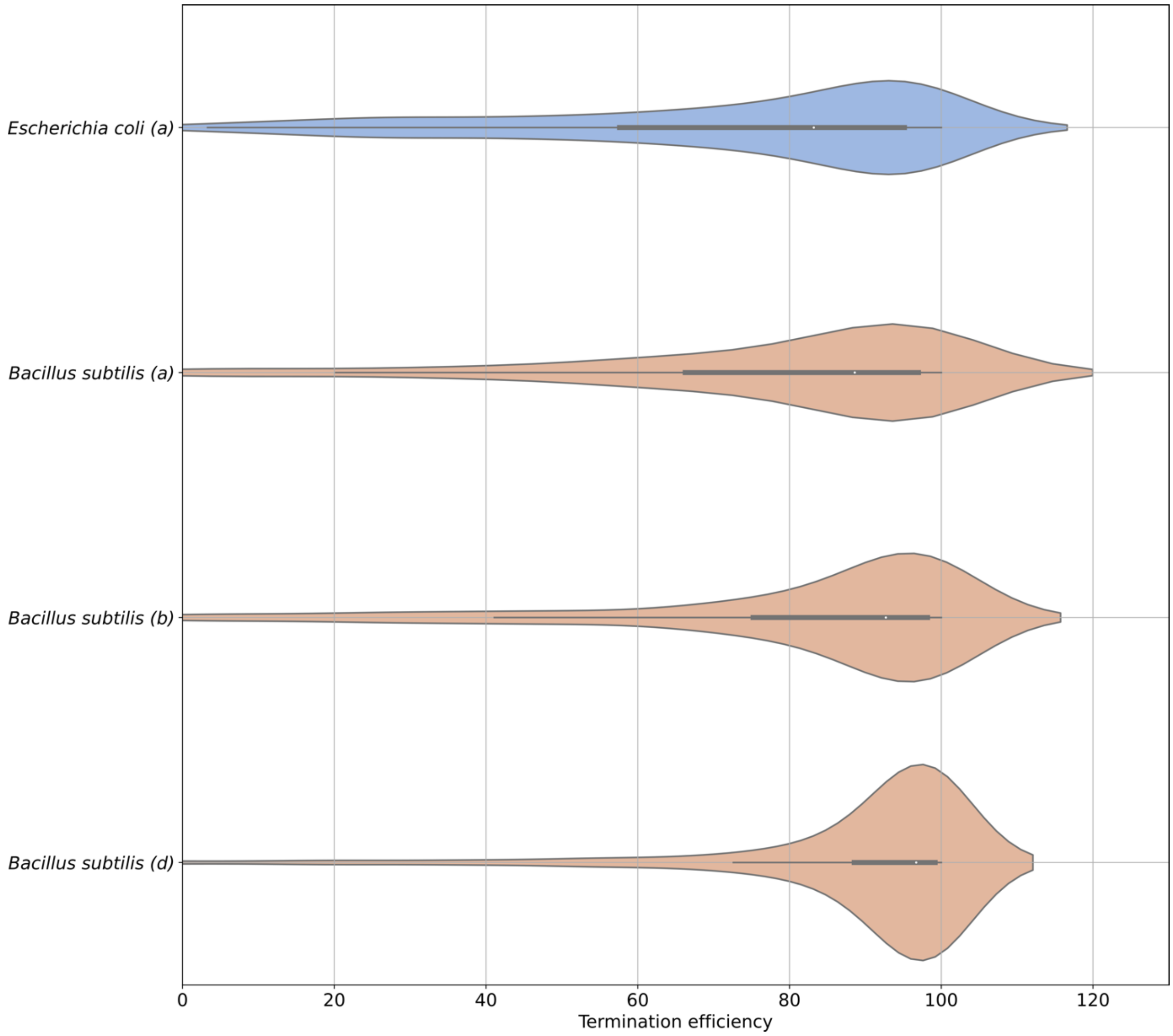


**Supplementary Figure 12.** Distribution of termination efficiencies calculated for all identified intrinsic terminators in each analyzed dataset prepared by the Term-seq protocol published by Mondal et al.

**Supplementary Table 1.** Description of Term-seq datasets used in the study. [Table provided as separate Excel file, available online]

**Supplementary Table 2.** The complete set of intrinsic terminators identified in the study. [Table provided as separate Excel file, available online]

**Supplementary Table 3.** Descriptive statistics calculated to characterize distributions of hairpin Minimum Free Energy (MFE) values of all identified intrinsic terminators in each dataset. Abbreviations: *nobs*, number of observations; *Q1*, first quartile; *Q3*, third quartile; *min*, minimum; *max*, maximum; *mean*, arithmetic mean; *variance*, unbiased variance (*nobs -1* was used as a denominator); *skewness*, distribution skewness with denominator equal to *nobs*; *kurtosis*, normalized kurtosis of the distribution (kurtosis = 0 for the normal distribution), no degrees of freedom were used for the calculations.

| | nobs | Q1 | median | Q3 | min | max | mean | variance | skewness | kurtosis |
|---|---|---|---|---|---|---|---|---|---|---|
| *Escherichia coli (a)* | 650 | -18.3 | -15.2 | -12.2 | -28.9 | -4.9 | -15.29 | 17.65 | -0.22 | -0.05 |
| *Escherichia coli (b)* | 904 | -18.6 | -15.2 | -12.08 | -28.5 | -3.9 | -15.36 | 18.74 | -0.22 | -0.44 |
| *Bacillus subtilis (a)* | 564 | -16.3 | -13.5 | -11.4 | -27.9 | -5.6 | -13.94 | 14.19 | -0.55 | 0.32 |
| *Bacillus subtilis (b)* | 1005 | -16.5 | -13.4 | -11.0 | -27.9 | -4.0 | -13.77 | 15.84 | -0.36 | 0.15 |
| *Bacillus subtilis (c)* | 867 | -16.5 | -13.7 | -11.5 | -27.9 | -4.1 | -13.99 | 14.36 | -0.43 | 0.25 |
| *Bacillus subtilis (d)* | 1053 | -16.4 | -13.4 | -11.2 | -27.9 | -4.0 | -13.8 | 15.17 | -0.39 | 0.21 |
| *Enterococcus faecalis* | 673 | -20.5 | -16.3 | -13.1 | -34.9 | -3.6 | -16.94 | 28.0 | -0.36 | -0.13 |
| *Listeria monocytogenes* | 754 | -17.0 | -14.1 | -11.7 | -26.2 | -3.3 | -14.23 | 15.25 | -0.12 | -0.09 |
| *Zymomonas mobilis* | 189 | -18.8 | -16.8 | -14.7 | -28.4 | -7.2 | -16.89 | 11.54 | -0.29 | 0.55 |
| *Synechocystis sp.* | 154 | -15.58 | -13.3 | -10.6 | -23.8 | -5.3 | -13.24 | 13.01 | -0.27 | -0.08 |
| *Streptomyces coelicolor* | 607 | -29.45 | -23.7 | -14.7 | -43.0 | -4.2 | -22.91 | 80.12 | -0.09 | -0.99 |
| *Streptomyces venezuelae* | 548 | -30.53 | -25.95 | -17.9 | -43.4 | -3.4 | -24.45 | 74.69 | 0.29 | -0.73 |
| *Streptomyces tsukubensis* | 499 | -31.65 | -25.3 | -16.3 | -43.0 | -3.2 | -24.15 | 84.63 | 0.2 | -1.03 |
| *Streptomyces griseus* | 701 | -33.1 | -25.9 | -15.8 | -44.7 | -6.1 | -24.91 | 94.04 | 0.1 | -1.18 |
| *Streptomyces lividans* | 471 | -28.65 | -23.6 | -15.45 | -42.1 | -7.4 | -22.68 | 68.33 | 0.03 | -0.92 |
| *Streptomyces clavuligerus* | 414 | -30.1 | -25.5 | -16.97 | -44.1 | -3.4 | -23.78 | 72.95 | 0.26 | -0.75 |
| *Streptomyces avermitilis* | 667 | -30.5 | -25.0 | -17.0 | -42.4 | -3.4 | -23.82 | 73.72 | 0.19 | -0.91 |

**Supplementary Table 4.** Results of the two-sided Mann-Whitney U rank test performed for each pairwise comparison of samples on distributions of multiple studied intrinsic terminators features. Statistic - Mann-Whitney U statistic corresponding to the sample A; P-value - associated p-value; FDR - P-value corrected for false discovery rate using the method of Benjamini/Hochberg; FDR < 0.05 - True if FDR < 0.05, False otherwise. [Table provided as separate Excel file, available online]

**Supplementary Table 5.** Descriptive statistics calculated to characterize distributions of hairpin scores calculated by the TransTermHP pipeline for all identified intrinsic terminators annotated in each dataset. Abbreviations: *nobs* – number of observations; *Q1* – first quartile; *Q3* – third quartile; *min* - minimum; *max* - maximum; *mean* - arithmetic mean; *variance* - unbiased variance, *nobs -1* is used as a denominator; *skewness* – distribution skewness with denominator equal to *nobs*; *kurtosis* - normalized kurtosis of the distribution (kurtosis = 0 for the normal distribution), no degrees of freedom are used for the calculations.

| | nobs | Q1 | median | Q3 | min | max | mean | variance | skewness | kurtosis |
|---|---|---|---|---|---|---|---|---|---|---|
| *Escherichia coli (a)* | 530 | -14.08 | -12.4 | -10.4 | -21.5 | -3.0 | -12.28 | 8.35 | -0.01 | 0.36 |
| *Escherichia coli (b)* | 661 | -14.0 | -12.2 | -10.3 | -21.5 | -3.0 | -12.13 | 8.85 | -0.02 | 0.16 |
| *Bacillus subtilis (a)* | 569 | -13.2 | -11.2 | -9.0 | -21.3 | -2.6 | -11.1 | 10.26 | -0.1 | 0.07 |
| *Bacillus subtilis (b)* | 1039 | -13.2 | -11.1 | -8.8 | -20.9 | -2.3 | -11.03 | 11.03 | -0.08 | -0.08 |
| *Bacillus subtilis (c)* | 885 | -13.3 | -11.3 | -9.1 | -22.4 | -2.4 | -11.28 | 10.77 | -0.09 | 0.14 |
| *Bacillus subtilis (d)* | 1080 | -13.2 | -11.2 | -8.9 | -21.3 | -2.4 | -11.11 | 10.8 | -0.08 | -0.04 |
| *Enterococcus faecalis* | 666 | -16.3 | -12.7 | -9.5 | -29.3 | -2.4 | -12.96 | 24.53 | -0.26 | -0.37 |
| *Listeria monocytogenes* | 797 | -12.9 | -10.4 | -8.1 | -24.6 | -2.2 | -10.59 | 12.96 | -0.27 | 0.01 |
| *Zymomonas mobilis* | 165 | -14.7 | -13.2 | -11.1 | -19.8 | -2.9 | -12.95 | 8.75 | 0.04 | 0.28 |
| *Synechocystis sp.* | 119 | -9.8 | -7.9 | -5.45 | -14.6 | -2.1 | -7.72 | 8.42 | -0.09 | -0.72 |
| *Streptomyces coelicolor* | 222 | -24.9 | -21.45 | -18.3 | -40.9 | -8.1 | -21.8 | 29.07 | -0.4 | 0.75 |
| *Streptomyces venezuelae* | 243 | -24.95 | -21.4 | -18.5 | -37.8 | -3.1 | -21.38 | 28.85 | 0.31 | 0.56 |
| *Streptomyces tsukubensis* | 219 | -24.95 | -21.4 | -17.8 | -37.4 | -5.6 | -21.41 | 31.81 | 0.04 | 0.14 |
| *Streptomyces griseus* | 280 | -27.5 | -22.95 | -19.2 | -42.7 | -7.4 | -23.47 | 31.9 | -0.11 | -0.01 |
| *Streptomyces lividans* | 195 | -23.75 | -20.8 | -17.9 | -32.4 | -8.1 | -20.91 | 21.74 | -0.01 | 0.27 |
| *Streptomyces clavuligerus* | 217 | -24.0 | -20.7 | -18.1 | -35.2 | -2.1 | -20.78 | 26.87 | 0.5 | 1.49 |
| *Streptomyces avermitilis* | 325 | -24.0 | -20.8 | -17.6 | -35.1 | -4.4 | -20.91 | 26.11 | 0.02 | -0.03 |

**Supplementary Table 6.** Descriptive statistics calculated to characterize distributions of hairpin lengths of all identified intrinsic terminators in each dataset. Hairpin length is defined as the number of nucleotides classified as a part of the hairpin loop or stem. Abbreviations: *nobs,* number of observations; *Q1*, first quartile; *Q3*, third quartile; *min*, minimum; *max*, maximum; *mean*, arithmetic mean; *variance*, unbiased variance (*nobs -1* was used as a denominator); *skewness*, distribution skewness with denominator equal to *nobs*; *kurtosis*, normalized kurtosis of the distribution (kurtosis = 0 for the normal distribution), no degrees of freedom were used for the calculations.

| | nobs | Q1 | median | Q3 | min | max | mean | variance | skewness | kurtosis |
|---|---|---|---|---|---|---|---|---|---|---|
| *Escherichia coli (a)* | 691 | 20.0 | 24.0 | 28.5 | 13 | 56 | 24.6 | 44.97 | 0.97 | 1.45 |
| *Escherichia coli (b)* | 957 | 20.0 | 25.0 | 30.0 | 13 | 51 | 25.47 | 45.97 | 0.6 | 0.08 |
| *Bacillus subtilis (a)* | 635 | 21.0 | 24.0 | 27.5 | 11 | 48 | 24.97 | 31.08 | 0.97 | 1.21 |
| *Bacillus subtilis (b)* | 1165 | 21.0 | 24.0 | 28.0 | 11 | 54 | 25.12 | 33.96 | 0.91 | 1.16 |
| *Bacillus subtilis (c)* | 984 | 21.0 | 24.0 | 28.0 | 11 | 49 | 24.97 | 30.97 | 1.0 | 1.43 |
| *Bacillus subtilis (d)* | 1214 | 21.0 | 24.0 | 28.0 | 11 | 54 | 25.1 | 34.09 | 1.02 | 1.66 |
| *Enterococcus faecalis* | 796 | 27.0 | 33.0 | 38.0 | 12 | 59 | 32.86 | 58.21 | 0.17 | -0.19 |
| *Listeria monocytogenes* | 862 | 24.0 | 28.0 | 33.0 | 11 | 57 | 28.66 | 48.13 | 0.76 | 0.74 |
| *Zymomonas mobilis* | 206 | 24.0 | 27.0 | 32.75 | 15 | 59 | 29.0 | 54.92 | 1.31 | 2.41 |
| *Synechocystis sp.* | 165 | 23.0 | 27.0 | 32.0 | 12 | 59 | 27.52 | 61.01 | 0.87 | 1.52 |
| *Streptomyces coelicolor* | 629 | 28.0 | 33.0 | 38.0 | 14 | 58 | 32.83 | 44.99 | 0.05 | 0.38 |
| *Streptomyces venezuelae* | 573 | 28.0 | 33.0 | 39.0 | 14 | 59 | 33.36 | 54.89 | 0.26 | 0.79 |
| *Streptomyces tsukubensis* | 520 | 29.0 | 34.0 | 39.0 | 16 | 57 | 33.84 | 49.51 | 0.0 | 0.34 |
| *Streptomyces griseus* | 737 | 29.0 | 34.0 | 38.0 | 14 | 57 | 33.4 | 44.89 | -0.1 | 0.62 |
| *Streptomyces lividans* | 495 | 28.0 | 32.0 | 37.0 | 16 | 58 | 32.38 | 46.49 | 0.12 | 0.28 |
| *Streptomyces clavuligerus* | 435 | 29.0 | 35.0 | 39.0 | 13 | 59 | 34.37 | 53.36 | 0.14 | 0.26 |
| *Streptomyces avermitilis* | 705 | 28.0 | 33.0 | 38.0 | 16 | 59 | 33.01 | 47.06 | 0.29 | 0.63 |

**Supplementary Table 7.** Descriptive statistics calculated to characterize distributions of hairpin loop lengths for all identified intrinsic terminators annotated in each experiment and species. Abbreviations: *nobs* – number of observations; *Q1* – first quartile; *Q3* – third quartile; *min* - minimum; *max* - maximum; *mean* - arithmetic mean; *variance* - unbiased variance, *nobs -1* is used as a denominator; *skewness* – distribution skewness with denominator equal to *nobs*; *kurtosis* - normalized kurtosis of the distribution (kurtosis = 0 for the normal distribution), no degrees of freedom are used for the calculations.

| | nobs | Q1 | median | Q3 | min | max | mean | variance | skewness | kurtosis |
|---|---|---|---|---|---|---|---|---|---|---|
| *Escherichia coli (a)* | 530 | 4.0 | 4.0 | 5.0 | 3 | 11 | 4.4 | 1.47 | 1.46 | 3.12 |
| *Escherichia coli (b)* | 661 | 4.0 | 4.0 | 5.0 | 3 | 11 | 4.38 | 1.53 | 1.51 | 3.24 |
| *Bacillus subtilis (a)* | 569 | 4.0 | 4.0 | 5.0 | 3 | 11 | 4.61 | 2.27 | 1.17 | 1.35 |
| *Bacillus subtilis (b)* | 1039 | 3.0 | 4.0 | 5.0 | 3 | 12 | 4.58 | 2.37 | 1.42 | 2.61 |
| *Bacillus subtilis (c)* | 885 | 3.0 | 4.0 | 5.0 | 3 | 12 | 4.59 | 2.29 | 1.29 | 2.12 |
| *Bacillus subtilis (d)* | 1080 | 3.0 | 4.0 | 5.0 | 3 | 12 | 4.59 | 2.4 | 1.4 | 2.56 |
| *Enterococcus faecalis* | 666 | 4.0 | 4.0 | 6.0 | 3 | 13 | 4.87 | 2.93 | 1.4 | 2.81 |
| *Listeria monocytogenes* | 797 | 4.0 | 4.0 | 6.0 | 3 | 12 | 4.74 | 2.48 | 1.29 | 2.26 |
| *Zymomonas mobilis* | 165 | 4.0 | 4.0 | 5.0 | 3 | 10 | 4.45 | 1.75 | 1.06 | 1.18 |
| *Synechocystis sp.* | 119 | 3.0 | 4.0 | 5.5 | 3 | 10 | 4.61 | 2.37 | 1.08 | 0.92 |
| *Streptomyces coelicolor* | 222 | 4.0 | 4.0 | 5.0 | 3 | 11 | 4.56 | 2.13 | 1.45 | 2.62 |
| *Streptomyces venezuelae* | 243 | 4.0 | 4.0 | 5.0 | 3 | 10 | 4.35 | 1.53 | 1.62 | 3.45 |
| *Streptomyces tsukubensis* | 219 | 4.0 | 4.0 | 5.0 | 3 | 12 | 4.55 | 2.35 | 1.67 | 3.78 |
| *Streptomyces griseus* | 280 | 4.0 | 4.0 | 5.0 | 3 | 13 | 4.55 | 2.31 | 2.03 | 6.1 |
| *Streptomyces lividans* | 195 | 4.0 | 4.0 | 5.0 | 3 | 9 | 4.5 | 1.78 | 1.25 | 1.59 |
| *Streptomyces clavuligerus* | 217 | 4.0 | 4.0 | 5.0 | 3 | 10 | 4.48 | 2.19 | 1.66 | 3.22 |
| *Streptomyces avermitilis* | 325 | 4.0 | 4.0 | 5.0 | 3 | 12 | 4.43 | 2.27 | 2.09 | 5.75 |

**Supplementary Table 8.** Descriptive statistics calculated to characterize distributions of tail scores of all identified intrinsic terminators in each dataset. Abbreviations: *nobs*, number of observations; *Q1*, first quartile; *Q3*, third quartile; *min*, minimum; *max*, maximum; *mean*, arithmetic mean; *variance*, unbiased variance (*nobs -1* was used as a denominator); *skewness*, distribution skewness with denominator equal to *nobs*; *kurtosis*, normalized kurtosis of the distribution (kurtosis = 0 for the normal distribution), no degrees of freedom were used for the calculations.

| | nobs | Q1 | median | Q3 | min | max | mean | variance | skewness | kurtosis |
|---|---|---|---|---|---|---|---|---|---|---|
| *Escherichia coli (a)* | 530 | -5.5 | -4.8 | -3.9 | -6.4 | -2.5 | -4.65 | 0.87 | 0.39 | -0.9 |
| *Escherichia coli (b)* | 661 | -5.4 | -4.7 | -3.8 | -6.4 | -2.5 | -4.58 | 0.91 | 0.28 | -1.04 |
| *Bacillus subtilis (a)* | 569 | -5.6 | -5.3 | -4.7 | -6.4 | -2.7 | -5.12 | 0.54 | 1.16 | 0.99 |
| *Bacillus subtilis (b)* | 1039 | -5.6 | -5.3 | -4.6 | -6.4 | -2.5 | -5.05 | 0.63 | 1.06 | 0.54 |
| *Bacillus subtilis (c)* | 885 | -5.6 | -5.3 | -4.7 | -7.1 | -2.6 | -5.11 | 0.56 | 1.13 | 0.97 |
| *Bacillus subtilis (d)* | 1080 | -5.6 | -5.3 | -4.6 | -6.4 | -2.5 | -5.04 | 0.63 | 1.05 | 0.51 |
| *Enterococcus faecalis* | 666 | -5.7 | -5.3 | -4.7 | -7.0 | -2.5 | -5.13 | 0.63 | 1.0 | 0.64 |
| *Listeria monocytogenes* | 797 | -5.7 | -5.5 | -5.1 | -6.5 | -2.9 | -5.31 | 0.37 | 1.37 | 2.27 |
| *Zymomonas mobilis* | 165 | -5.6 | -5.3 | -4.4 | -6.2 | -2.7 | -4.95 | 0.81 | 0.97 | -0.18 |
| *Synechocystis sp.* | 119 | -5.6 | -5.3 | -4.9 | -6.2 | -3.1 | -5.18 | 0.37 | 1.06 | 0.86 |
| *Streptomyces coelicolor* | 222 | -3.8 | -3.5 | -3.1 | -5.5 | -2.5 | -3.52 | 0.35 | -0.52 | 0.22 |
| *Streptomyces venezuelae* | 243 | -4.1 | -3.6 | -3.1 | -5.5 | -2.5 | -3.59 | 0.43 | -0.57 | -0.09 |
| *Streptomyces tsukubensis* | 219 | -4.0 | -3.5 | -3.1 | -5.8 | -2.5 | -3.56 | 0.47 | -0.86 | 0.58 |
| *Streptomyces griseus* | 280 | -3.7 | -3.4 | -2.9 | -5.5 | -2.5 | -3.43 | 0.38 | -0.77 | 0.37 |
| *Streptomyces lividans* | 195 | -3.8 | -3.5 | -3.1 | -5.6 | -2.5 | -3.55 | 0.4 | -0.65 | 0.35 |
| *Streptomyces clavuligerus* | 217 | -3.9 | -3.5 | -3.1 | -5.8 | -2.5 | -3.57 | 0.5 | -1.01 | 0.7 |
| *Streptomyces avermitilis* | 325 | -3.9 | -3.4 | -3.1 | -5.9 | -2.5 | -3.54 | 0.44 | -0.76 | 0.38 |

**Supplementary Table 9.** Descriptive statistics calculated to characterize distributions of the distances between the POT and the nearest upstream gene located on the same strand in each dataset. Abbreviations: *nobs*, number of observations; *Q1*, first quartile; *Q3*, third quartile; *min*, minimum; *max*, maximum; *mean*, arithmetic mean; *variance*, unbiased variance (*nobs -1* was used as a denominator); *skewness*, distribution skewness with denominator equal to *nobs*; *kurtosis*, normalized kurtosis of the distribution (kurtosis = 0 for the normal distribution), no degrees of freedom were used for the calculations.

| | nobs | Q1 | median | Q3 | min | max | mean | variance | skewness | kurtosis |
|---|---|---|---|---|---|---|---|---|---|---|
| *Escherichia coli (a)* | 691 | 38.0 | 48.0 | 69.5 | 2.0 | 199.0 | 62.76 | 1492.51 | 1.83 | 2.65 |
| *Escherichia coli (b)* | 957 | 38.0 | 48.0 | 73.0 | 1.0 | 200.0 | 63.9 | 1640.91 | 1.7 | 2.12 |
| *Bacillus subtilis (a)* | 635 | 33.0 | 38.0 | 46.0 | 6.0 | 193.0 | 46.28 | 814.43 | 2.93 | 9.56 |
| *Bacillus subtilis (b)* | 1165 | 32.0 | 38.0 | 45.0 | 8.0 | 196.0 | 45.03 | 797.73 | 2.82 | 9.19 |
| *Bacillus subtilis (c)* | 984 | 32.0 | 38.0 | 44.0 | 8.0 | 196.0 | 44.13 | 750.1 | 3.03 | 10.56 |
| *Bacillus subtilis (d)* | 1214 | 31.0 | 38.0 | 45.0 | 8.0 | 196.0 | 44.75 | 824.32 | 2.81 | 9.02 |
| *Enterococcus faecalis* | 796 | 42.0 | 51.0 | 71.0 | 1.0 | 200.0 | 62.76 | 1216.6 | 1.77 | 3.12 |
| *Listeria monocytogenes* | 862 | 37.0 | 41.0 | 49.0 | 1.0 | 199.0 | 47.73 | 676.87 | 3.29 | 13.16 |
| *Zymomonas mobilis* | 206 | 49.0 | 57.0 | 70.0 | 4.0 | 198.0 | 66.34 | 918.05 | 1.96 | 4.22 |
| *Synechocystis sp.* | 165 | 68.0 | 91.0 | 126.0 | 1.0 | 200.0 | 98.7 | 1568.61 | 0.52 | -0.02 |
| *Streptomyces coelicolor* | 629 | 56.0 | 76.0 | 118.0 | 4.0 | 200.0 | 90.29 | 2121.29 | 0.78 | -0.42 |
| *Streptomyces venezuelae* | 573 | 53.0 | 67.0 | 100.0 | 1.0 | 200.0 | 80.66 | 1658.47 | 1.08 | 0.61 |
| *Streptomyces tsukubensis* | 520 | 52.0 | 67.0 | 100.25 | 1.0 | 198.0 | 79.47 | 1719.81 | 0.99 | 0.33 |
| *Streptomyces griseus* | 737 | 56.0 | 69.0 | 97.0 | 1.0 | 200.0 | 80.42 | 1494.59 | 1.04 | 0.69 |
| *Streptomyces lividans* | 495 | 52.5 | 66.0 | 99.0 | 7.0 | 200.0 | 81.11 | 1794.79 | 1.12 | 0.6 |
| *Streptomyces clavuligerus* | 435 | 54.0 | 70.0 | 111.0 | 2.0 | 200.0 | 84.89 | 1943.18 | 0.93 | 0.1 |
| *Streptomyces avermitilis* | 705 | 52.0 | 67.0 | 99.0 | 1.0 | 200.0 | 79.9 | 1637.05 | 1.1 | 0.63 |

**Supplementary Table 10.** Descriptive statistics calculated to characterize distributions of distances between the POT and the nearest downstream gene located on the same strand for all identified intrinsic terminators annotated in each experiment and species. Abbreviations: *nobs* – number of observations; *Q1* – first quartile; *Q3* – third quartile; *min* - minimum; *max* - maximum; *mean* - arithmetic mean; *variance* - unbiased variance, *nobs -1* is used as a denominator; *skewness* – distribution skewness with denominator equal to *nobs*; *kurtosis* - normalized kurtosis of the distribution (kurtosis = 0 for the normal distribution), no degrees of freedom are used for the calculations.

| | nobs | Q1 | median | Q3 | min | max | mean | variance | skewness | kurtosis |
|---|---|---|---|---|---|---|---|---|---|---|
| *Escherichia coli (a)* | 691 | 82.0 | 264.0 | 1633.0 | 3.0 | 40597.0 | 1469.56 | 8905260.06 | 5.59 | 52.15 |
| *Escherichia coli (b)* | 957 | 105.0 | 375.0 | 1812.0 | 2.0 | 40598.0 | 1704.31 | 10829139.36 | 4.69 | 33.97 |
| *Bacillus subtilis (a)* | 635 | 82.0 | 211.0 | 1305.0 | 11.0 | 58833.0 | 1671.35 | 20704051.81 | 7.27 | 72.94 |
| *Bacillus subtilis (b)* | 1165 | 95.0 | 353.0 | 1697.0 | 11.0 | 76739.0 | 1833.31 | 21231131.38 | 8.1 | 98.32 |
| *Bacillus subtilis (c)* | 984 | 92.75 | 325.5 | 1683.0 | 11.0 | 76739.0 | 1882.39 | 23045489.27 | 8.14 | 97.11 |
| *Bacillus subtilis (d)* | 1214 | 95.0 | 351.5 | 1759.0 | 11.0 | 76739.0 | 1864.88 | 21777302.75 | 7.76 | 90.79 |
| *Enterococcus faecalis* | 794 | 95.0 | 282.0 | 2079.0 | 2.0 | 30260.0 | 2155.74 | 20104345.95 | 3.56 | 14.12 |
| *Listeria monocytogenes* | 862 | 80.0 | 234.0 | 1802.5 | 7.0 | 77755.0 | 2303.26 | 34549442.77 | 6.29 | 56.09 |
| *Zymomonas mobilis* | 206 | 137.25 | 425.0 | 2058.25 | 5.0 | 14228.0 | 1707.09 | 6762451.13 | 2.18 | 4.72 |
| *Synechocystis sp.* | 165 | 80.0 | 177.0 | 1325.0 | 1.0 | 8612.0 | 1125.82 | 3665028.21 | 2.35 | 5.08 |
| *Streptomyces coelicolor* | 629 | 142.0 | 732.0 | 2302.0 | 1.0 | 31398.0 | 1985.13 | 10383444.1 | 3.4 | 17.33 |
| *Streptomyces venezuelae* | 573 | 84.0 | 450.0 | 1834.0 | 4.0 | 31480.0 | 1664.19 | 9388157.6 | 3.96 | 22.85 |
| *Streptomyces tsukubensis* | 519 | 148.0 | 558.0 | 2330.5 | 6.0 | 71482.0 | 2139.07 | 21523262.55 | 7.8 | 98.54 |
| *Streptomyces griseus* | 737 | 113.0 | 562.0 | 1918.0 | 1.0 | 68511.0 | 1907.85 | 18322104.81 | 8.62 | 111.62 |
| *Streptomyces lividans* | 495 | 100.5 | 357.0 | 1784.0 | 4.0 | 42700.0 | 1671.99 | 11546888.55 | 5.53 | 48.7 |
| *Streptomyces clavuligerus* | 435 | 141.0 | 484.0 | 1967.0 | 4.0 | 23628.0 | 1724.57 | 7674920.0 | 2.91 | 12.09 |
| *Streptomyces avermitilis* | 705 | 108.0 | 534.0 | 1927.0 | 1.0 | 78901.0 | 1751.21 | 16617207.04 | 11.07 | 186.94 |

**Supplementary Table 11.** statistics calculated to characterize distributions of distances between the hairpin base and the POT for intrinsic terminators identified by the TERMITe in each dataset. Abbreviations: *nobs*, number of observations; *Q1*, first quartile; *Q3*, third quartile; *min*, minimum; *max*, maximum; *mean*, arithmetic mean; *variance*, unbiased variance (*nobs -1* was used as a denominator); *skewness*, distribution skewness with denominator equal to *nobs*; *kurtosis*, normalized kurtosis of the distribution (kurtosis = 0 for the normal distribution), no degrees of freedom were used for the calculations.

| | nobs | Q1 | median | Q3 | min | max | mean | variance | skewness | kurtosis |
|---|---|---|---|---|---|---|---|---|---|---|
| *Escherichia coli (a)* | 691 | 3.0 | 5.0 | 6.0 | 0.0 | 10.0 | 4.24 | 3.89 | -0.39 | -0.32 |
| *Escherichia coli (b)* | 957 | 3.0 | 5.0 | 6.0 | 0.0 | 10.0 | 4.6 | 4.77 | -0.33 | -0.12 |
| *Bacillus subtilis (a)* | 635 | 5.0 | 6.0 | 7.0 | 0.0 | 10.0 | 5.42 | 3.62 | -1.14 | 1.0 |
| *Bacillus subtilis (b)* | 1165 | 5.0 | 6.0 | 7.0 | 0.0 | 10.0 | 5.33 | 4.17 | -0.98 | 0.47 |
| *Bacillus subtilis (c)* | 984 | 5.0 | 6.0 | 7.0 | 0.0 | 10.0 | 5.58 | 3.79 | -1.16 | 0.9 |
| *Bacillus subtilis (d)* | 1214 | 5.0 | 6.0 | 7.0 | 0.0 | 10.0 | 5.46 | 3.78 | -1.1 | 0.73 |
| *Enterococcus faecalis* | 796 | 4.0 | 6.0 | 7.0 | 0.0 | 10.0 | 5.17 | 5.42 | -0.71 | -0.35 |
| *Listeria monocytogenes* | 862 | 5.0 | 6.0 | 7.0 | 0.0 | 10.0 | 5.82 | 3.27 | -1.37 | 1.85 |
| *Zymomonas mobilis* | 206 | 3.0 | 5.0 | 6.0 | 0.0 | 10.0 | 4.48 | 4.88 | -0.47 | -0.3 |
| *Synechocystis sp.* | 165 | 5.0 | 6.0 | 7.0 | 0.0 | 10.0 | 5.49 | 3.63 | -0.49 | 1.25 |
| *Streptomyces coelicolor* | 629 | 2.0 | 4.0 | 6.0 | 0.0 | 10.0 | 4.26 | 7.71 | 0.29 | -0.7 |
| *Streptomyces venezuelae* | 573 | 2.0 | 4.0 | 6.0 | 0.0 | 10.0 | 3.89 | 6.99 | 0.19 | -0.8 |
| *Streptomyces tsukubensis* | 520 | 2.0 | 4.0 | 5.0 | 0.0 | 10.0 | 3.71 | 6.65 | 0.36 | -0.55 |
| *Streptomyces griseus* | 737 | 2.0 | 4.0 | 6.0 | 0.0 | 10.0 | 4.13 | 7.65 | 0.17 | -0.83 |
| *Streptomyces lividans* | 495 | 2.0 | 4.0 | 6.0 | 0.0 | 10.0 | 4.13 | 5.96 | 0.09 | -0.64 |
| *Streptomyces clavuligerus* | 435 | 2.0 | 5.0 | 7.0 | 0.0 | 10.0 | 4.91 | 8.12 | -0.05 | -1.05 |
| *Streptomyces avermitilis* | 705 | 2.0 | 4.0 | 6.0 | 0.0 | 10.0 | 4.0 | 6.68 | 0.26 | -0.55 |

**Supplementary Table 12.** Descriptive statistics calculated to characterize distributions of termination efficiencies calculated for all identified intrinsic terminators in each experiment prepared by the protocol published by Mondal et al. (Mondal et al., 2016). Abbreviations: *nobs* – number of observations; *Q1* – first quartile; *Q3* – third quartile; *min* - minimum; *max* - maximum; *mean* - arithmetic mean; *variance* - unbiased variance, *nobs -1* is used as a denominator; *skewness* - skewness with denominator equal to *nobs*; *kurtosis* - normalized kurtosis (kurtosis = 0 for the normal distribution), no degrees of freedom are used for the calculations.

| | nobs | Q1 | median | Q3 | min | max | mean | variance | skewness | kurtosis |
|---|---|---|---|---|---|---|---|---|---|---|
| *Escherichia coli (a)* | 611 | 57.6 | 83.2 | 95.15 | -78.5 | 100.0 | 72.75 | 764.92 | -1.19 | 1.09 |
| *Bacillus subtilis (a)* | 571 | 66.25 | 88.6 | 97.0 | -382.4 | 100.0 | 76.67 | 1106.9 | -5.51 | 63.24 |
| *Bacillus subtilis (b)* | 1165 | 75.2 | 92.7 | 98.2 | -80.9 | 100.0 | 80.89 | 686.98 | -1.94 | 3.66 |
| *Bacillus subtilis (d)* | 1214 | 88.52 | 96.7 | 99.2 | -35.1 | 100.0 | 88.69 | 405.28 | -2.92 | 9.05 |
